# Quantitative Analysis of Transcriptome Dynamics Provides Novel Insights into Developmental State Transitions

**DOI:** 10.1101/2022.03.10.483850

**Authors:** Kristin Johnson, Simon Freedman, Rosemary Braun, Carole LaBonne

**Author notes:** Corresponding author: Carole LaBonne.

## Abstract

During embryogenesis, the developmental potential of initially pluripotent cells becomes progressively restricted as they transit to lineage restricted states. The pluripotent cells of *Xenopus* blastula-stage embryos are an ideal system in which to study cell state transitions during developmental decision-making, as gene expression dynamics can be followed at high temporal resolution. Here we use transcriptomics to interrogate the process by which pluripotent cells transit to four different lineage-restricted states: neural progenitors, epidermis, endoderm and ventral mesoderm, providing quantitative insights into the dynamics of Waddington’s landscape. Our findings shed light on why the neural progenitor state is the default lineage state for pluripotent cells, and uncover novel components of lineage-specific gene regulation. These data reveal an unexpected overlap in the transcriptional responses to BMP4/7 and activin signaling, and provide mechanistic insight into how the timing of signaling inputs such as BMP are temporally controlled to ensure correct lineage decisions. Together these analyses provide quantitative insights into the logic and dynamics of developmental decision making in early embryos.

## Introduction

How a single cell ultimately gives rise to a patterned, complex organism is a fundamental question in biology. Embryonic development can be generalized as a process of progressive restriction of cellular potential. In vertebrates, the zygote is totipotent, but by blastula stages the three primary germ layers, ectoderm, mesoderm and endoderm, have been specified. The fates of cells within these germ layers then become progressively restricted to single differentiated cell types characteristic of that germ layer. Conrad Waddington famously depicted this process as a topological landscape (Waddington 1957). In his model, a ball positioned at the top of the landscape represents a cell with all developmental pathways open to it. As the ball progresses down the landscape, the paths it takes dictate which lineage states will remain accessible. Waddington noted that the valleys or channels of the landscape arise from the interactions between genes and from their interactions with the cell’s environment.

At blastula stages vertebrate embryos possess a transient population of pluripotent cells which, like the fertilized egg, occupy a position at the top of Waddington’s landscape. These cells-inner cell mass cells in mammals and naïve animal pole cells in amphibians-can give rise to the derivative cell types of all three germ layers and as such can recapitulate the path to different lineage states including the relevant gene regulatory network (GRN) topology and dynamics. Studies using explants of pluripotent cells from *Xenopus* blastulae (so called “animal caps”) have been central to our current understanding of the signals and transcriptional responses that direct these stem cells toward specific lineage states (Snape et al. 1987, Ariizumi and Asashima 2001, Ariizumi et al. 2009, Ariizumi et al. 2017, Satou-Kobayashi et al. 2021). Some of these signals emanate from the blastopore lip, or the Spemann-Mangold organizer, and help to direct formation and patterning of the primary germ layers (Spemann & Mangold 1924, Harland & Gerhart 1997, Niehrs 2004). Exit from the pluripotent state also coincides with the loss of expression of many maternally provided pluripotency transcripts, such as *pou5f3.3* and *foxi2* (Whitfield et al. 1995, Lef et al. 1994, Paraiso et al. 2020). As cells exit pluripotency, their potential becomes progressively restricted until their fate becomes specified and then determined.

Inner animal pole cells in *Xenopus* are fated to give rise to ectodermal derivatives, and when explanted from the embryo will form epidermis absent additional signals or instructions (Jones and Woodland 1986). In situ these cells will give rise to both epidermal and neural progenitor cells, as well as neural crest and cranial placodes, under the direction of signals secreted from the organizer. Absent these signals, animal pole cells are directed by endogenous BMP signaling to become epidermis (Wilson et al. 1997). BMP2, 4, and 7 have all been shown to be potent epidermal inducers (Wilson and Hemmati-Brivanlou 1995, Suzuki et al. 1997), and BMP4/7 heterodimers have been identified as the most physiologically relevant ligands in early embryos (Little and Mullins 2009). The binding of these ligands to type I and II BMP receptors results in phosphorylation of smad1/5/8 and the translocation of these phospho-smads to the nucleus together with smad4, resulting in transcription of target genes including *epidermal keratin* (*EpK*) and *dlx3* (Kretzschmar et al. 1997, Macias Silva et al. 1998, Shi and Massague 2003, Jonas et al. 1985, Dirksen et al. 1994).

Although isolated animal pole cells will transit to an epidermal state absent additional instructions, it has been proposed that the default state of these initially pluripotent cells is a neural progenitor state (Hemmati-Brivanlou and Melton 1997). This model arose from the findings that neural “inducing” factors secreted by the organizer, including noggin, chordin, follistatin and cerberus, function as BMP antagonists (Lamb et al. 1993, Sasai et al. 1995, Piccolo et al. 1996, Piccolo et al. 1999, Iemura at al. 1998, Munoz-Sanjuan & Brivanlou 2002). The “neural default” model has not been without controversy, however. In chick embryos the expression patterns of BMPs and their antagonists do not fully fit the neural default model (Streit et al. 1998), and misexpression of BMP antagonists does not induce neural progenitors, nor does ectopic expression of BMP inhibit neural plate formation in this system (Streit and Stern 1999a, Streit and Stern 1999b, Stern 2005). By contrast, *Xenopus* animal pole explants exposed to BMP antagonists such as noggin adopt a neural progenitor state and express neural genes including Sox2/3 and *Otx1/2*. These transcription factors, and their homologs, play an important role in specification of the central nervous system (CNS) in vertebrates (Collignon et al. 1996, Mizuseki et al. 1998, Penzel et al. 2003, Pannese et al, 1995, Kablar et al. 1996, Andreazzoli et al. 1997, Plouhinec et al. 2017).

Explants of *Xenopus* pluripotent blastula cells have also played a central role in determining the signals that control formation of the other embryonic germ layers, mesoderm and endoderm (Asashima et al. 1990a,b, Smith et al. 1990, Henry, et al. 1996). Nodal, activin and Vg-1, ligands of the other branch of the TGF-beta family, have been shown to act as morphogens, with low levels of signaling inducing mesoderm and high levels inducing endoderm (Smith et al. 1990, Hematti-Brivanlou & Melton 1992, Gurdon, et al. 1994, McDowell et al 1997, McDowell and Gurdon 1999, Agius et al. 2000, Dale et al. 1993, Thomsen and Melton 1993, Kessler and Melton 1996). Treatment of these cells with exogenous activin mimics the activity of nodal/Vg-1 (Jones et al. 1995) promoting phosphorylation of receptor smads 2/3 (Reissmann et al. 2001, Kumar et al. 2001). The maternal transcription factor VegT also plays a key role in mesendoderm formation. VegT can induce endoderm both cell-autonomously and via its induction of TGF-beta signaling, whereas its mesoderm inducing activity is indirect via TGF-beta signaling, helping to distinguish these two lineage states (Zhang et al. 1998, Kimelman & Griffin 1998, Clements et al. 1999). High levels of activin/nodal signaling induce endoderm as evidenced by expression of key factors *Sox17* and *endodermin* (Hudson et al. 1997, Sasai et al. 1996) whereas lower doses induce both dorsal and ventral mesoderm (Sokol and Melton 1992, Green et al. 1992). BMP4/7 heterodimers also possess mesoderm inducing activity, but it is limited to ventral but not dorsal mesoderm (Nishimatsu and Thomsen 1998, Hemmati-Brivanlou and Thomsen 1995, Graff. et al 1994). BMP4 homodimers were initially shown to have mesoderm-inducing activity, weakly inducing genes such as the t-box transcription factor *brachyury (t)* and the hox gene *evx1* (Koster et al. 1991, Suzuki et al. 1997, Smith et al. 1991, Altaba et al. 1989). Subsequently BMP7 was found to have weak mesoderm inducing and ventralizing activities (Dick et al 2000, Schmid et al 2000). More recently, it has been shown that BMP7 acts primarily as a heterodimer (Kim et al. 2019). While the ability of BMP4/7 to induce ventral mesoderm has been established, much still remains to be learned about the signaling dynamics and outputs of BMP4/7 versus activin-mediated regulation of cell state transitions during germ layer formation.

*Xenopus* animal pole explants are an ideal system to probe the dynamics of developmental decision-making during lineage restriction. These cells can be cultured in a simple salt solution, and will undergo lineage restriction on the same time scale as if they had remained in vivo. Given appropriate signals, animal pole cells can be directed to any lineage progenitor state within a time frame of approximately seven hours. Analogous experiments in cultured mouse or human embryonic stem cells (ESCs) would take more than a week in culture, and unlike ESCs *Xenopus* cells do not need to be artificially retained in a pluripotent state that may be distinct from the transient pluripotency that exists in vivo. The unique features of the *Xenopus* system allow initially pluripotent cells to be followed at high temporal resolution as they progress to lineage restriction, providing insights into the dynamics of developmental decision making in early vertebrate embryos.

Here we develop an experimental platform and quantitative framework in which pluripotent cells explanted from blastula stage *Xenopus* embryos can be used to study the transit of these cells to four different lineage states - epidermis, neural progenitor, endoderm and ventral mesoderm - by following changes in the transcriptome at six time points during this seven-hour process. These data provide quantitative insights into the dynamics of Waddington’s landscape. Our findings shed light on why a neural progenitor state is the default lineage state for pluripotent cells, uncover novel components of lineage specific GRNs, and provide insights into essential control of the timing of signaling inputs such as BMP for proper lineage decisions. These time-resolved data sets will serve as an important resource for future studies of developmental decision making in early vertebrate embryos.

## Results

### Naïve Animal Pole Cells from *Xenopus* Blastula can be Programmed to any Lineage State

To allow interrogation of transcriptome changes at high temporal resolution as pluripotent cells become lineage restricted, we established a highly regimented protocol for collecting synchronous populations of blastula explants across six points on the path towards lineage restriction. Late blastula stage explants (Nieuwkoop and Faber stage 9) were designated time zero and represent the pluripotent state atop Waddington’s landscape (Fig 1a). In addition to time zero, explants were collected at 75, 150, 225, 315 and 435 minutes after stage 9, (Nieuwkoop and Faber stages 10, 10.5, 11, 12, 13), confirmed by the stage of sibling embryos (Nieuwkoop and Faber 1994). Stage 10.5 is the onset of gastrulation as marked by the presence of the dorsal blastopore lip (Fig 1b) whereas stages 11 and 12 are mid and late gastrulae respectively. Stage 13, the neural plate stage, is the stage by which the developmental potential of embryonic cells has been restricted to a single lineage state, with the notable exception of neural crest cells (Prasad et al. 2012). Explants cultured without additional instructive cues transit to an epidermal state (Fig. 1C). To follow cells as they transited to an endodermal state, explants were treated with 160ng/uL of activin at stage 9. Similarly, to follow transit to a neural progenitor state, explants were exposed to 100ng/uL of the BMP antagonist noggin. Treatment with 20ng/uL BMP 4/7 was used to induce a ventral mesoderm state and to allow a comparison of the transcriptional responses to the two different branches of TGF-beta signaling. RNA was isolated from explants at each time interval and used to generate illumina libraries for transcriptome analysis.

**Figure 1.**
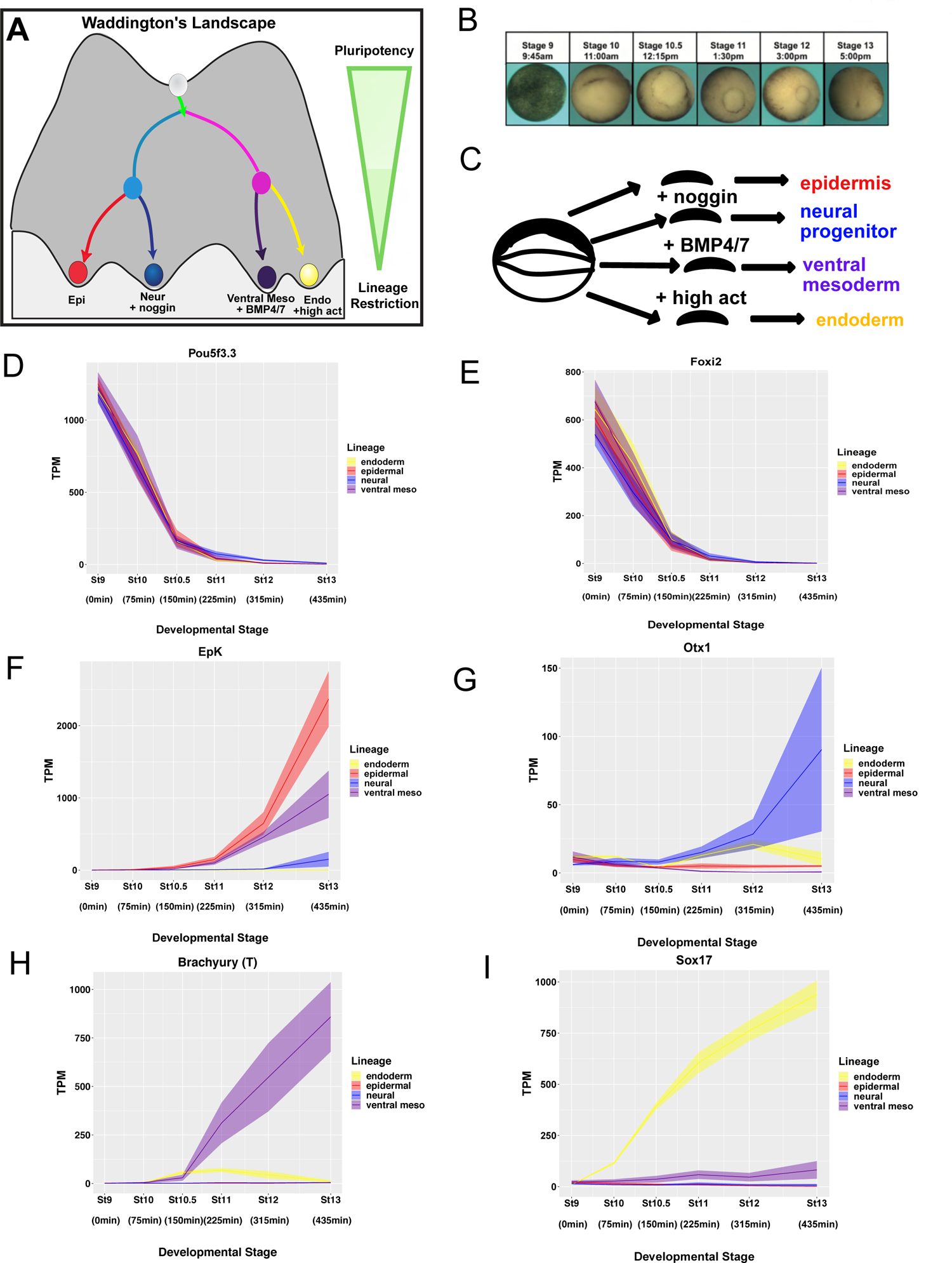
Xenopus blastula explants can be reprogrammed to any lineage state. (A) Schematic of Waddington’s Landscape portraying lineage specification process. (B) Embryos stage 9,10,10.5,11,12, and 13 used to confirm developmental stages of blastula explants. (C) Schematic showing signaling molecules used to direct 4 distinct lineage transitions. (D-I) RNA Seq TPM expression over time of (D) maternally provided pluripotency marker *Pou5f3.3,* (E) maternally provided *Foxi2*, (F) epidermal marker *EpK*, (G) neural marker *Otx1,* (H) mesoderm marker *Brachyury(T)* (I) endoderm marker *Sox17* (H) Graphs are sums of S+L allele. Width of lines represents SEM of three biological replicates.

The transcript dynamics of maternally provided pluripotency factors *Pou5f3.3* and *Foxi2* across three biological replicates for all four lineage transitions (twelve independent experiments) demonstrates that this pipeline produced highly quantitative and highly reproducible data with minimal technical error (Fig. 1D,E). *Pou5f3.3* and *Foxi2* are representative of a class of 119 genes whose expression decreases monotonically by at least 25-fold between stage 9 and stage 13 (Fig. S1 A) inclusive of maternally provided transcripts, many of which characterize the pluripotent state (Collart et al. 2014, Gentsch et al. 2019). Normalized RNA read count data, quantified as transcripts per million (TPM), confirmed generation of each of the expected lineage states with *EpK* and *dlx3* validating establishment of an epidermal state (Fig. 1F and S1B), *Otx1* and *Otx2* validating transit to a neural state (Fig. 1G and S1C). *Brachyury(T)* and *Evx1* (Fig. 1H and S1D), and *Sox17* and *Endodermin* (Fig. 1I, S1E) validated the mesoderm and endoderm data sets respectively. Global analysis of transcription factors expressed at each stage revealed that the majority are expressed in all four lineages, although their expression levels or timing may vary between them. For example, *Otx1* is unique to the endodermal lineage at stage 10 but to the neural lineage at stage 13 (Fig. S2 A-E). This analysis also identified transcription factors that at a given stage were expressed in only a single lineage, and interestingly these genes were overrepresented in the endoderm lineage.

### PCA and Time Series Analysis Reveal Novel Lineage-Specific Dynamics

Together the transcriptomes of the four state transitions each across six time points yield 72 observations in a 45,661-dimensional gene expression space. Global insights into such high dimensional data require methods for dimensionality reduction. Principal Component Analysis (PCA) can provide key insights into the genes contributing most significantly to the variance between lineage states and developmental stages. We first used PCA to analyze each lineage individually, plotting the first two principal components against developmental time (Fig. 2A). For all four lineages the primary principal component (PC1) was found to be largely monotonic over time, suggesting that the majority of the gene expression variance is contributed by genes changing unidirectionally, such as pluripotency genes being turned off or lineage-specific genes being activated.

**Figure 2.**
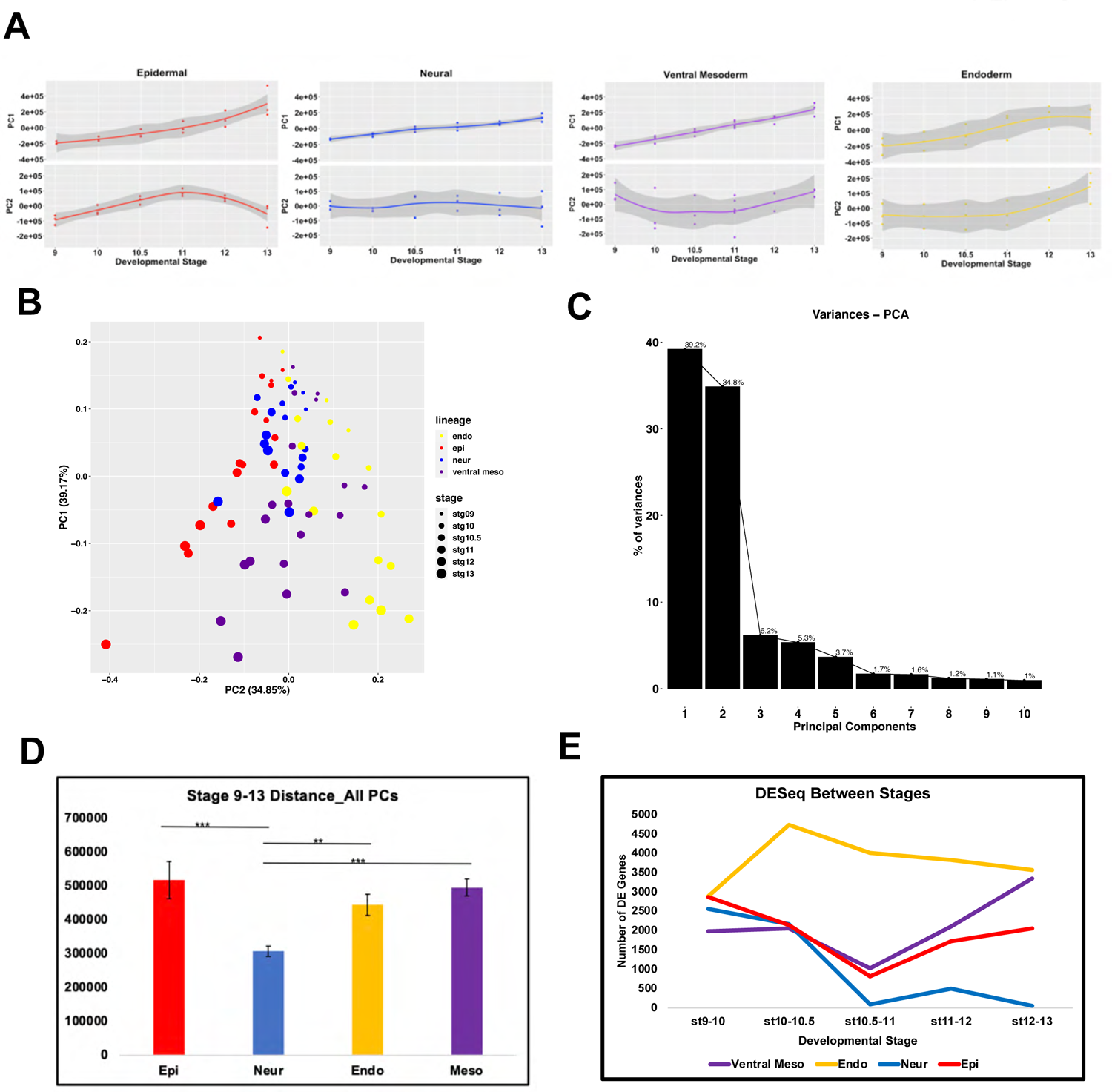
PCA and Time Series Analysis Reveal Novel Lineage-Specific Dynamics. (A) PCA for each individual lineage with the coordinates of PC1 and PC2 for each lineage plotted against developmental time. (B) PCA performed on all four lineages simultaneously, with plot showing clustering of all lineages for PC1 vs PC2. (C) Scree plot of the variance explained by the top10 principal components for PCA done on all lineages. (D) Distances from stage 9 to 13 for all PCs for each lineage, error bars are SEM of all 9 stage 9-13 distances for each lineage, (\*\*\**P*<0.005, **P<.01). (E) Number of genes differentially expressed between successive stages in each lineage, *p*_adj_ ≤ 0.05.

Interestingly, when PC2 was plotted for each of the lineages it was found to also exhibit temporal dynamics. The profiles for epidermis, ventral mesoderm and endoderm suggest that genes exhibiting expression peaks at intermediate time points make a significant contribution to the variance, and potentially to the state transitions themselves. By contrast, PC2 of the neural lineage shows no association with time, suggesting that the temporal dynamics in the neural lineage are primarily linear in time. This raises the possibility that the transition to a neural progenitor state follows a simpler trajectory than that of the other three lineages (Fig. 2A).

To gain further insights into these state transitions, PCA was carried out on all four lineages in concert. In this analysis, PC1 and PC2 together explain 75% of the variance across these data. (Fig. 2B, C). The distribution of these data across the PC1 axis corelates with developmental time, with the earliest (St.9) samples clustering at the top of the plot and the later samples progressively more distal (Fig. 2B). When examined in this context, the neural trajectory is striking as it extends a shorter distance along this axis, with the neural replicates for stages 11, 12 and 13 clustering closer to the stage 10.5 replicates for the other lineages. To quantify this and ensure this is not an artifact of dimension reduction, the distance between stages 9 and 13 in the full gene expression space was calculated. We find that here too the neural lineage moves the shortest distance (Fig. 2D). Whereas PC1 correlates with developmental time, PC2 appears to distinguish the different lineage states, which after stage 10.5 show very distinct trajectories. Interestingly, endoderm and epidermis lie furthest from each other along PC2 with mesoderm lying between those states. Thus, PC2 captures the intermediate nature of the ventral mesoderm state which shares GRN features with endoderm but, like epidermis, is BMP-driven (Fig. 2B).

We next used differential expression analysis (DESeq2) to gain insights into the dynamics of gene expression changes across the four state transitions. Plotting the number of genes differentially expressed between successive developmental stages reveals that the number of genes whose expression changes significantly during these state transitions is relatively modest (Fig. 2E). For example, between two and three thousand genes are differentially expressed between stages 9 and 10 in each lineage, which represents four to six percent of the transcriptome. Between stages 10 and 10.5, the gene expression changes in the endodermal lineage are strikingly different from those of the other lineages. The number of differentially expressed genes increases almost 60% between these stages in the endoderm, whereas there is a significant decrease in differentially expressed genes in the epidermal and neural lineages and little change in the mesoderm. While there is a gradual decrease in differentially expressed genes in the endodermal lineage between stages 10.5 and 13, this state transition continues to exhibit the most dynamic changes in gene expression compared to the other lineages. After a reaching a minimum between stages 10.5 and 11 both the epidermal and ventral mesodermal lineages exhibit increasing numbers of differentially expressed genes. By stages 12-13 the endoderm and ventral mesoderm exhibit comparable gene expression dynamics. Interestingly, almost a quarter of the genes changing in the endodermal and ventral mesodermal trajectories between stages 12 and 13 are shared between these lineages, likely reflecting the overlapping landscape of the combined mesendoderm GRN (Fig S3I-J). In contrast to the other three lineages, the neural trajectory exhibits very few differentially expressed genes after stage 11, providing additional evidence that this lineage reaches an early equilibrium (Fig. 2E, S3G-J).

### Gene Expression Dynamics Provide Novel Insights into the Neural Default State

The above analyses suggested that the neural progenitor state follows a simpler trajectory than that of the other three lineages. Comparison of gene expression dynamics during transit to an epidermal versus a neural state reveals that through stage 10.5 these two lineages share a remarkably similar trajectory (Fig. 3A). For each, the number of genes differentially expressed between successive stages decreases, the number of genes changing is highly similar, and there is significant overlap in the genes exhibiting differential expression. Between stages 9 and 10, for example, 60% of the genes differentially expressed in the neural linage are also differentially expressed in the epidermal lineage and that is true of 53% of genes differentially expressed between stages 10 and 10.5 (Fig. 3B). The majority of shared genes exhibit decreasing expression during these stages, in part reflecting the downregulation of pluripotency genes. Nevertheless, of the 1161 genes whose expression increases in the neural trajectory between stages 9 and 10, more than half also show increased expression in the epidermal trajectory.

**Figure 3.**
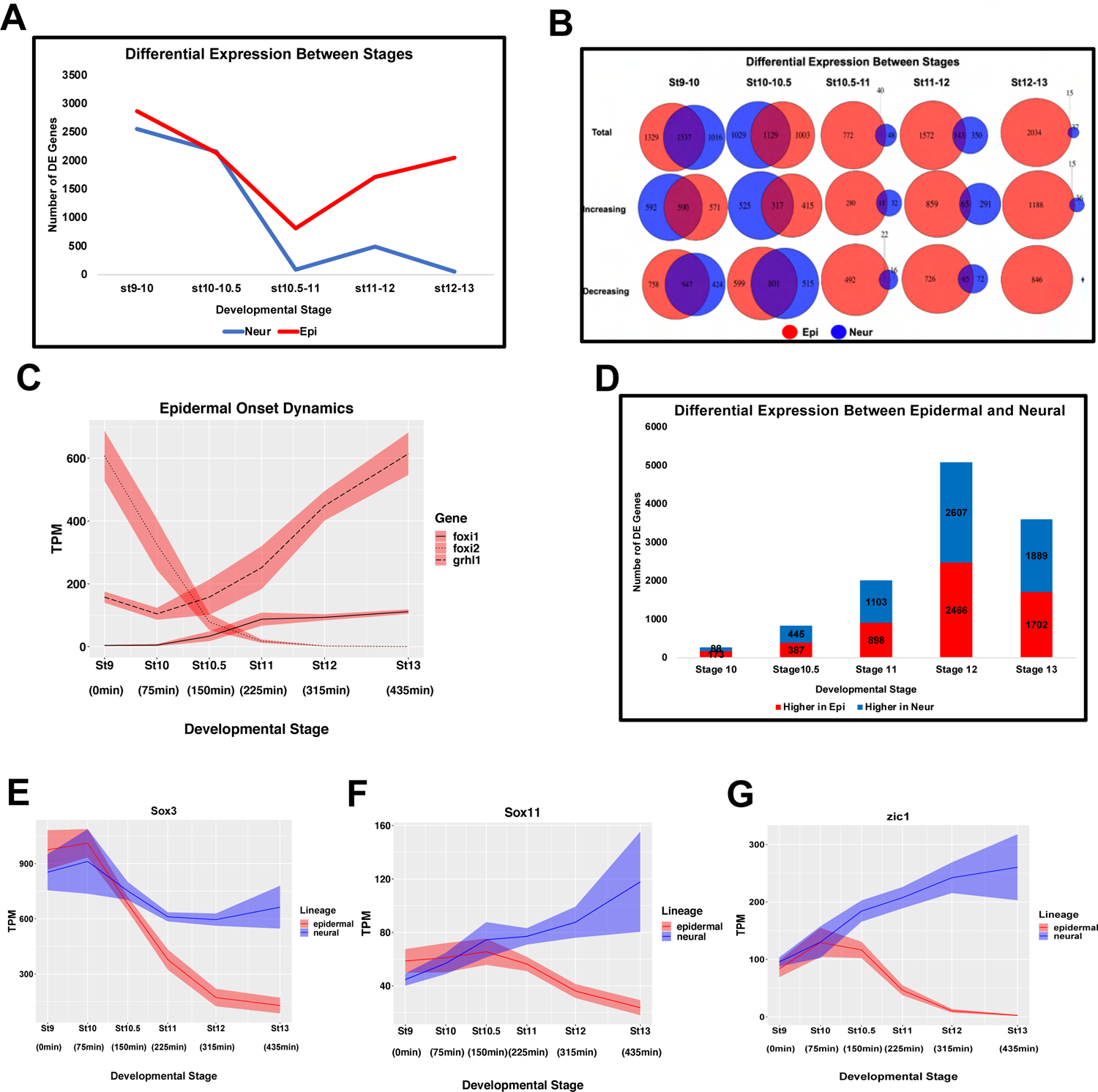
Gene Expression Dynamics Provide Novel Insights into the Neural Default State. (A) Number of differentially expressed genes between successive developmental stages of the epidermal and neural lineage, *p*_adj_ ≤ 0.05 (B) Venn Diagrams for the total number of DE genes between stages for the epidermal and neural lineages, as well as the number of genes increasing and decreasing over time. (C) TPM of *foxi1*, *foxi2* and *grhl1*, revealing epidermal onset dynamics. Graph shows sum of S+L allele. Width of the line represents SEM of three biological replicates. (D) Number of differentially expressed genes between lineages at each developmental stage (*p*_adj_ ≤ 0.05). (E-G) Graphs of the TPM of two pluripotency markers maintained in the neural lineage (E) *Sox3*, (F) *Sox11*, and (G) *zic1*. Graphs are sums of S+L allele. Width of the line represents SEM of three biological replicates.

Between stages 10.5 and 11, which corresponds to early gastrulation, there is a striking divergence of the epidermal and neural lineages. During these stages, gene expression dynamics largely cease in the neural lineage; fewer than 100 genes are differentially expressed between stages 10.5 and 11, and only 52 genes are differentially expressed between stages 12 and 13. By contrast, in the epidermal lineage temporal changes in gene expression begins to sharply increase and the overlap of these genes with those changing in the neural lineage is minimal (Fig. 3A,B). Of all genes differentially expressed between stages 12 and 13 in these two stage transitions, less than 1% are differentially expressed in both the neural and epidermal lineages, mainly because the neural lineage has ceased to change.

Interestingly, the increasing gene expression dynamics that characterizes the epidermal lineage at stage 11 coincides with a loss of enrichment for neural GO terms (Fig. S4 A). This suggests that pluripotent blastula cells possess neural-like features that begin to be lost around the onset of gastrulation as cells transit to an epidermal state, but are retained and reinforced in neural progenitor cells.

The observed gene expression dynamics and GO term enrichment together point to the onset of gastrulation as a critical point on the landscape topology of early developmental decision making. To further explore this, we examined the expression of *foxi1* and *grhl1* which are key upstream components of the GRN mediating the formation of epidermis (Mir et al. 2007; Tao et al. 2005).

Expression of *foxi1* has been shown to be activated by the pluripotency factor *foxi2* (Cha et al. 2012). Examining the expression dynamics of these three genes across the epidermal trajectory reveals a sharp increase in the expression of *grhl1* that correlates with rapidly extinguishing expression of *foxi2* and the onset of gastrulation (stage 10.5) (Fig. 3C). As the onset of gastrulation is also when gene expression dynamics virtually cease in the neural trajectory, this suggests attractor dynamics that favor a neural progenitor state and a network structure that requires cells to be actively propelled toward an alternative, in this case epidermal, state.

To further examine the genes that distinguish the neural and epidermal states we analyzed the genes differentially expressed between these states at each time point on their trajectories. At stage 10 the two lineages remain strikingly similar, with only 261 genes differentially expressed (Fig. 3D). The number of differentially expressed genes increases by more than 500% between stages 10.5 and 12, driven almost entirely by gene expression dynamics in the epidermal lineage. Interestingly, 13 of the top 20 most differentially expressed genes at stage 10.5 are known BMP responsive genes and all but three, *jun.L/S* and *actc1.S,* are more highly expressed in the epidermal lineage (Fig. S4 B). Consistent with this, beginning at stage10.5 genes associated with the TGF-beta pathway in the KEGG database show enrichment in the epidermal lineage (Fig. S4 C). As stage 10.5 represents the time when the trajectories of the neural and epidermal lineages diverge after neural reaches early equilibrium, this enrichment is consistent with a model where BMP signals actively propel cells away from a neural well and onto the path toward an epidermal state. Together, the early equilibrium reached by the neural lineage, combined with the neural features of the pluripotent state, help explain why neural is the default state following exit from pluripotency. This is further evidenced by the expression dynamics of genes that play important roles in both pluripotent cells and neural progenitors such as *Sox3, Sox11* and *Zic1* (Penzel et al. 2003, Hyodo-Miura et al. 2002; Mizuseki et al. 1998, Nakata et al. 1998). All three of these genes retain or increase their expression in the neural lineage but are rapidly down-regulated in the epidermal lineage after stage 10.5 (Fig. 3E-G).

### Robust BMP Signaling is Initiated in Explants Around the Onset of Gastrulation

Consistent with a model where BMP signals actively propel cells away from the neural state, phosphorylation of BMP R-Smads is first detected in animal pole explants at stage 10.5 (Fig. 4A). The translocation of pSmad1/5/8 to the nucleus drives expression of BMP responsive genes, and its timing correlates with the divergence of the epidermal lineage from neural lineage. To gain further insights the timing of BMP responsiveness we examined whether genes differentially expressed at successive developmental stages showed over-representation of genes associated with BMP signaling (Kanehisa and Goto 2000) using the DESeq2 Wald test. We computed the significance at which these BMP associated genes comprised a larger fraction than would be expected by random chance via the hypergeometric *p*-value (Virtanen et al. 2020). The greatest divergence in overrepresentation between the epidermal and neural lineages was seen between stages 10.5 and 11, which was also the maxima for overrepresentation in the epidermal data (Fig. 4B). Consistent with this finding, *Ventx2.1* and *Id3*, which are both BMP targets genes exhibit expression maxima in the epidermal lineage and minima in the neural lineages at these stages (Fig. 4C, D) (Onichtchouk et al. 1996, Hollnagel et al. 1999). The expression of *Id3* across these state transitions is particularly noteworthy for its opposite intermediate non-monotonic dynamics at successive developmental stages despite comparable expression at the start and end of these lineage trajectories.

**Figure 4.**
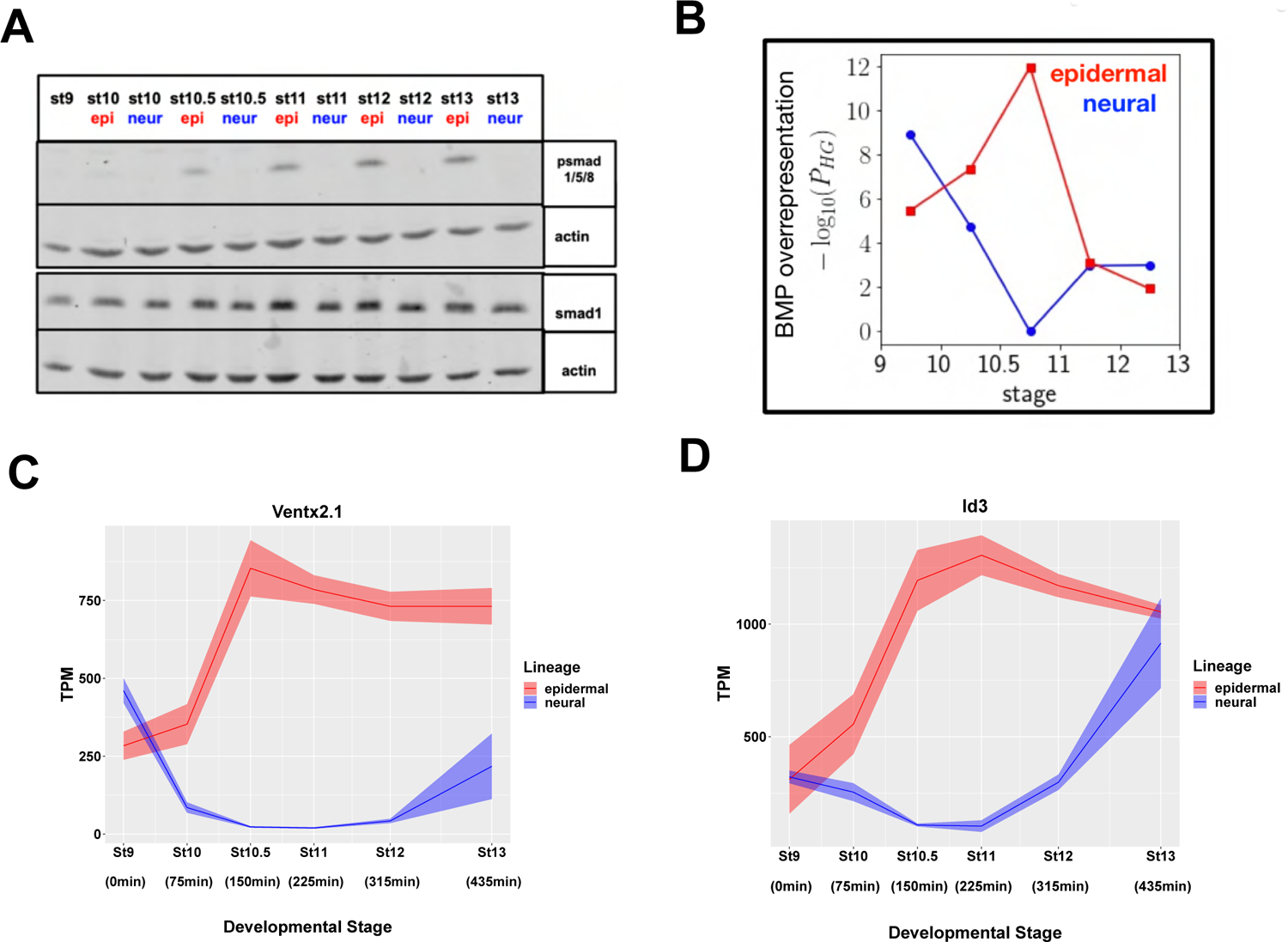
Robust BMP Signaling is initiated in Explants Around the Onset of Gastrulation. (A) Western blot analysis of lysates of developing epidermal (WT) and neural (20uM K02288) explants for psmad1/5/8 and smad1 with actin loading control. (B) Significance of BMP overrepresentation (hypergeometric p-value) in temporally differentially expressed genes (C-D) Graphs of BMP responsive genes in epidermal and neural lineages (C) *Ventx2.1* (D) *Id3*. Graphs are sums of S+L allele. Width of the line represents SEM of three biological replicates.

### Early Response to Activin and BMP4/7 Reveals Unexpected Overlap

While BMP signaling plays a central role in instructing pluripotent cells to form epidermis, it is the other branch of the TGF-β that directs mesendodermal fates. Members of the Activin/ Nodal/Vg1/GDF1/ TGF-beta subfamily act via pSmad2/3 to activate mesodermal and endodermal target genes including *foxh1*, *eomes*, *mixer*, *tcf3* (also known as *e2a*) and *tp53* (Chen at al. 1996, Ryan et al. 1996, Henry and Melton 1998, Rashbass et al 1992, Wills and Baker 2015, Cordenonsi et al. 2003.). While recent work has suggested that Vg1-Nodal heterodimers mediate this process endogenously (Montague and Schier 2017), activin has long been used for efficient mesendoderm induction in ex-vivo assays (Smith et al. 1990, Green et al. 1992, Hemmati-Brivanlou et al. 1992). Like activin, BMP4/7 heterodimers have been shown to be potent mesoderm inducers at physiological concentrations. However, in contrast to activin/nodal, which can induce ventral mesoderm, dorsal mesoderm and endoderm in a concentration dependent manner, BMP4/7 has been reported to induce only ventral mesodermal (Suzuki 1997, Nishimatsu and Thomsen GH 1998)). We focused on BMP4/7-mediated mesoderm induction for this analysis because the response to activin/nodal is a spectrum with no clear threshold cleanly distinguishing an endodermal versus mesodermal response. An additional advantage of examining BMP4/7-mediated mesoderm induction is that it provides an opportunity to directly compare the signaling dynamics and transcriptional responses of the two branches of TGF-beta signaling in the same quantitative framework.

Treatment with activin at stage 9 leads to robust signaling at stage 10 as evidenced by Western detection of phosphorylated-Smad2(p-Smad2) (Fig. S5 A) and robust induction of *Sox17* beginning at stage 10 (Fig 1I). Interestingly, transit to an endodermal state is distinguished from the other lineage transitions by its unique transcriptome dynamics. It is the only lineage in which there is a large increase in differentially expressed genes between stage 9 and stage 10.5, the onset of gastrulation, after which the number of differentially expressed genes decreases (Fig. 2D, 5A). By contrast, treatment of stage 9 explants with BMP4/7 does not elicit an immediate increase in differentially expressed genes, distinguishing the responses to the two different arms of TGF-beta signaling between successive developmental stages (Fig. 5A). Instead, the number of genes that are differentially expressed between stages 9-10 and 10-10.5 following BMP4/7 treatment remains fairly constant, before decreasing between stages 10.5 and 11. Intriguingly, however, the genes differentially expressed between these early stages in response to activin or BMP4/7 significantly overlap. For example, approximately 48% of genes differentially expressed between stages 9 and 10 in response to BMP4/7 are also differentially expressed between those stages in response to activin as are 71% of genes differentially expressed in response to BMP4/7 between stages 10 and 10.5 (Fig. 5B). These findings were unexpected as Smad1/5/8 and Smad2/3 generally regulate distinct target genes (Massagué and Wotton, 2000, Wardle and Smith, 2006).

**Figure 5.**
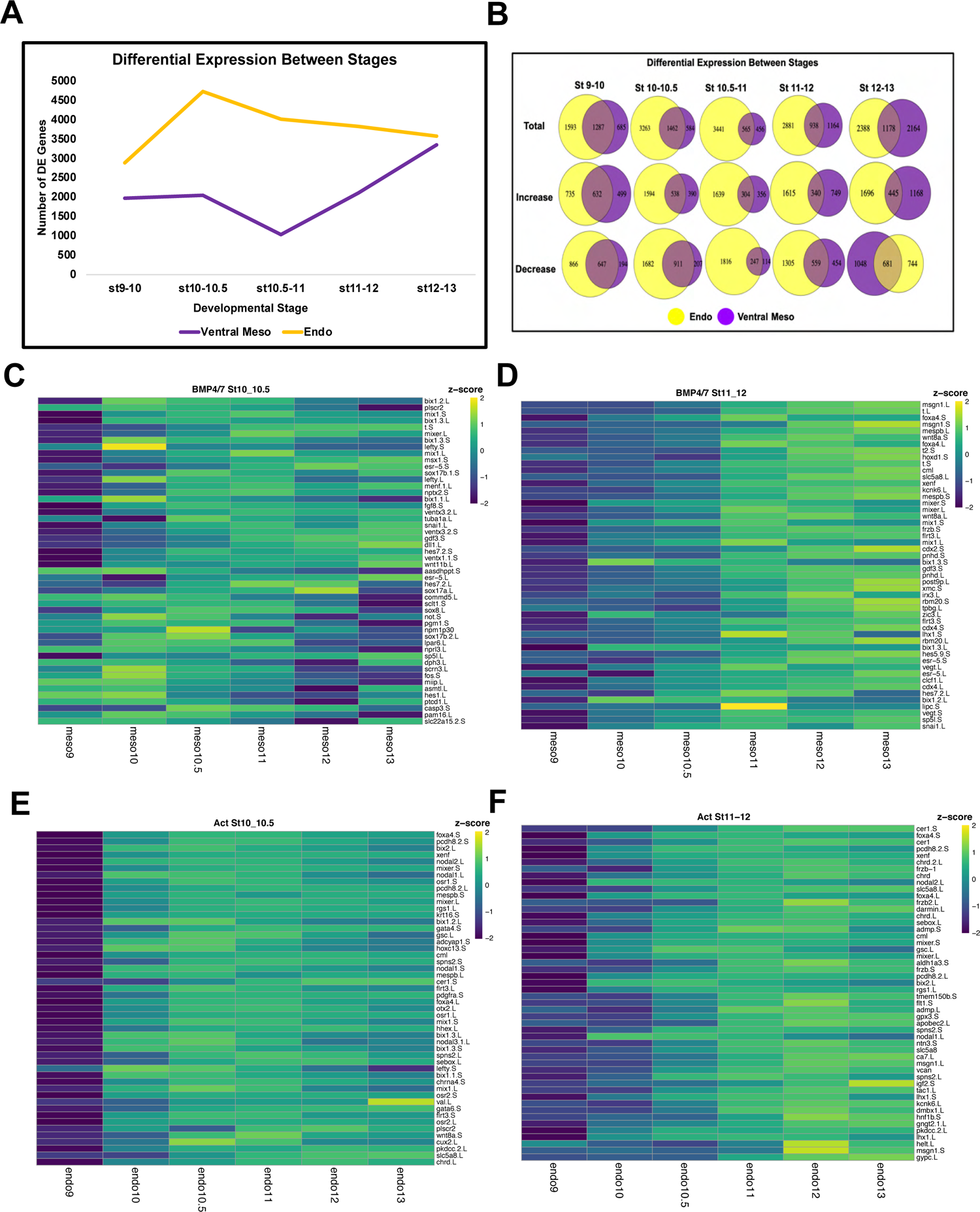
Early Response to Activin and BMP4/7 Reveals Unexpected Overlap. (A) Number of differentially expressed genes between successive developmental stages of the endoderm and ventral mesoderm lineage, *p*_adj_ ≤ 0.05 (B) Venn Diagrams for the total number of DE genes between stages for the epidermal and neural lineages, as well as the number of genes increasing and decreasing over time. (C-F) Heatmaps of the top 50 genes differentially increased in response to (C) BMP4/7 at stage 10 and/or 10.5 (D) BMP4/7 at stage 11 and/or 12 (E) Activin at stage 10 and/or 10.5 (F) Activin at stage 11 and/or 12. All genes in heatmaps are ranked by Log2FC of differential expression of either mesoderm (BMP4/7) or endoderm (Activin) compared to the wildtype epidermal lineage. Only genes increasing in response to BMP4/7 or Activin with a minimum expression of 10TPM at the relevant stages and higher expression at the relevant stage than at stage 9 were included in the heatmap. Colors represent *z*-scores of TPM.

To further explore the unexpected overlap in transcriptional responses to the two different classes of TGF-beta ligands we used DESeq2 to examine the genes that exhibit the largest log2 fold change compared to untreated explants in response to BMP4/7 treatment. Figure 5C shows the top 50 genes exhibiting the largest expression increase at stages 10 or 10.5 in response to BMP4/7 treatment relative to untreated explants. This gene set, which captures the immediate response to this ligand, includes previously characterized BMP target genes such as *msx1* and *ventx1* (Suzuki et al. 1997, Rastegar et al. 1999) as well as the pan-mesodermal gene *brachyury(t)* (Smith et al. 1991). Unexpectedly, it also includes a number of dorsal mesoderm/endoderm genes that have been characterized as targets of Activin/Nodal signaling including *bix1, mix1, mixer* and *sox17* (Tada et al. 1998, Rosa 1989, Chen et al 1996, Hudson et al. 1997). Notably, activation of these genes occurs absent activation of pSmad2/3 (Fig. S5 A). Using *z*-score scaling to visualize the expression dynamics of early responding genes across the time series revealed that genes generally associated with activin/nodal expression, including *bix1* and *mix1*, displayed non-monotonic dynamics with expression peaks at intermediate stages, whereas the expression of ventral and pan-mesodermal genes increased monotonically. (Figure 5C).

After reaching a minima between stages 10.5 and 11, the number of genes displaying dynamic expression changes in response to BMP4/7 greatly increased (Fig. 5A). The genes exhibiting the largest log2 fold change at stages 11 or 12 were therefore similarly examined. This gene set is more enriched for ventral mesoderm associated genes than the initially responding genes, suggesting that the activin-like response to BMP4/7 is transitory and that ventral mesoderm character is stabilized secondarily. This is consistent with a role for BMP signaling in actively ventralizing mesoderm and other tissues (Schmidt et al. 1995).

Interestingly, while treatment with both activin and BMP4/7 was initiated at stage 9, activin-mediated phosphorylation of smad2 was transient (Fig. S5 A) whereas BMP4/7-mediated phosphorylation of smad1/5/8 persisted through to lineage restriction at stage 13 (Fig 7A), likely contributing to the distinct gene expression dynamics in these two lineages. For example, the genes exhibiting the largest log2 fold change compared to untreated explants at stages 10, 10.5 or 11, 12 in response to activin treatment are enriched for those whose maximal expression occurs at intermediate stages of the lineage trajectory (Fig. 5E,F). Importantly, while the genes activated as an early response to activin or BMP4/7 show significant overlap (Fig.5B), the genes activated by these two classes of TGF-beta ligands nevertheless show significant differential expression with respect to one another (Fig. S5 B). Over 1000 genes are differentially expressed between these trajectories as early as stage 10, and the number of genes differentially expressed between these lineages continues to increase over developmental time.

Interestingly, analysis of KEGG pathway enrichment in these differentially expressed genes reveals that genes that are significantly higher in the endoderm lineage show enrichment for the TGF-beta pathway at stages 10 and 10.5, whereas genes that are significantly higher in the ventral mesoderm lineage are enriched for the TGF-beta pathway at stages 11 and 12 (Fig. S5 C).

### Time Series Data Provides Novel Insights into Mesendoderm GRN

The mesendoderm gene regulatory network (GRN) has been extensively studied and has yielded a significant “parts list” of genes that make significant contributions to the formation of these lineages (Charney et al. 2017, Jansen et al 2022), however the ordering of the GRN components has lacked the temporal resolution that our time series data can provide. Accordingly, we examined the expression of forty-one validated mesendoderm GRN components across both the endoderm and ventral mesoderm lineage trajectories (Fig. 6A, B). Interestingly, many of these genes display non-monotonic expression, with their expression peaking at early time points before decreasing. These dynamics are particularly striking in the endoderm, and demonstrate that many of these GRN components respond to activin rapidly and robustly, but transiently. Indeed, expression of twenty-one of these genes is induced in the endoderm by stage 10, which is 75 minutes after ligand exposure (Fig. 6A, B). Interestingly, sixteen of these genes are also activated by BMP4/7 by stage10, albeit less robustly, indicating that they are immediate responders to both classes of TGF-beta ligands. The activin-induced expression of seven of GRN components, including Bix and Nodal family genes, *vegt, eomes*, *mix1* and *snai1*, peaks by stage 10.5 and then declines. By contrast, a second group of GRN factors, including Sox17 and gata2/6-related genes, *osr2* and *pitx2* exhibit sustained expression through stage 13. A third group of GRN components, including *hnf1b*, *foxa2* and *gata5*, are not robustly turned on until mid- to late-gastrula stages (Fig 6A, B). The expression of *ventx* genes, known for their strong ventralizing activity, in response to high levels of activin signaling was unexpected, and correlates with the down-regulation of Nodal and Bix family factors. Surprisingly, *ventx2.1* and *ventx2.2*, which are among the first genes to respond robustly to BMP4/7, were induced as or more strongly by activin, albeit with different temporal dynamics. Subsequently GRN factors more closely associated with mesoderm, including *t* (*brachyury*), *wnt8*, and *msx1/2,* distinguish the BMP4/7 response from the activin response. As with the activin response, these BMP4/7 responding genes show distinct patterns of temporal dynamics (Fig. 6C).

**Figure 6.**
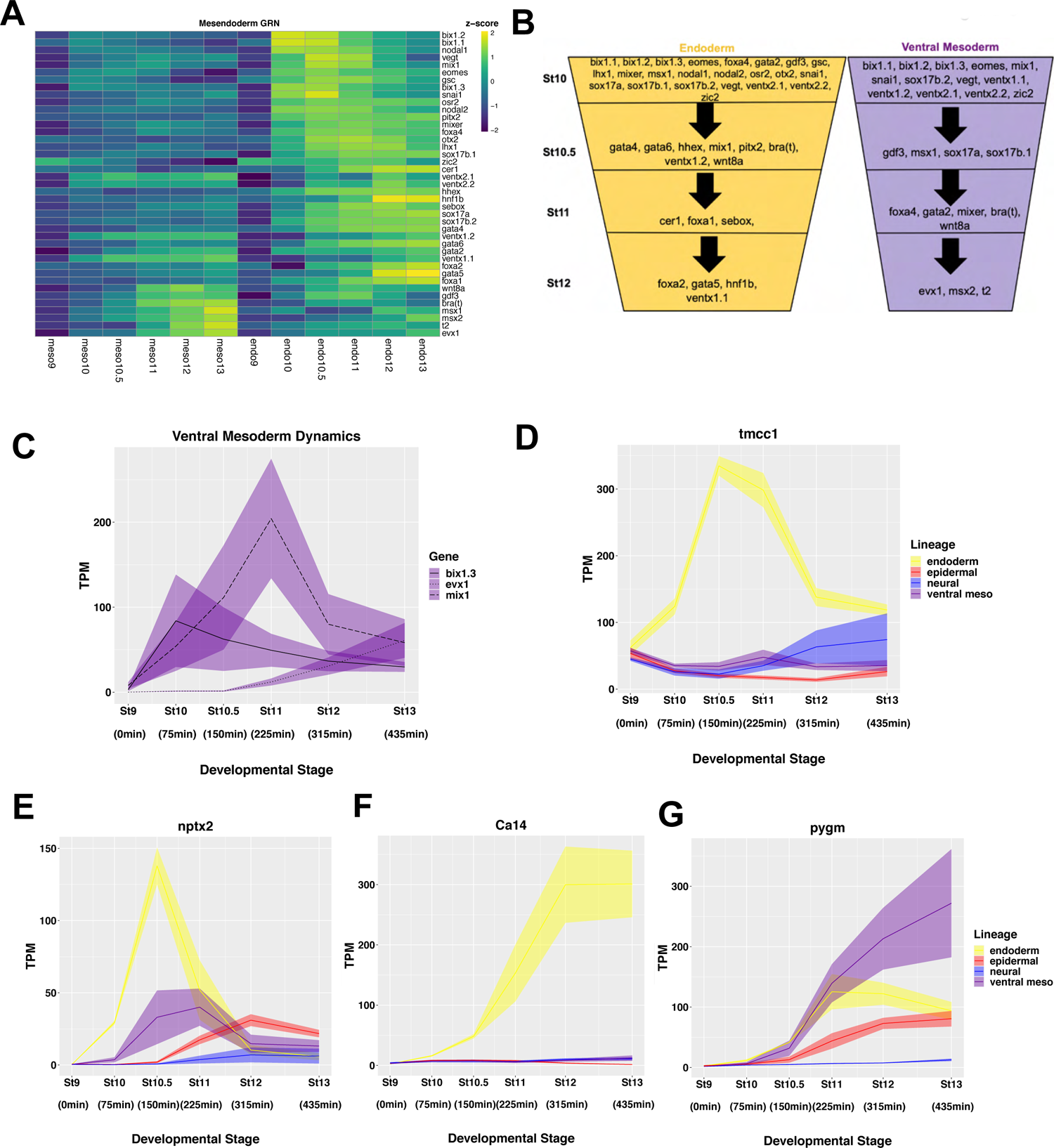
Time Series Data Provides Novel Insights into Mesendoderm GRN. (A) Heatmaps of genes in the published mesendoderm GRN across time in the endoderm and ventral mesoderm lineage with colors representing *z*-score of TPM expression across both lineages. (Charney et al. 2017) (B) Schematic of the timing of genes from published Mesendoderm GRN in both the endoderm and ventral mesoderm lineage, as defined by expression of at least 30 TPM in the L and S allele combined for the average of three biological replicates. (C) TPM of *bix1.3*, *evx1*, and *mix1* revealing mesoderm onset dynamics. (D-G) TPM of genes proposed as novel mesendoderm GRN members based on DESeq2 and Limma analysis (A) *tmcc1* (B) *nptx2* (C) *lrp4* (D) *pygm*. Graphs are sums of S+L allele. Width of the line represents SEM of three biological replicates.

Given the distinct dynamics displayed by known mesendoderm GRN factors, we tested for differential linear and quadratic dynamics in the BMP4/7 and activin responses using the R package limma (Law et al. 2014). This analysis identified genes displaying expression dynamics that position them as candidates for novel components the mesendoderm GRN. For example, one noted pattern was genes expressed rapidly and robustly in response to activin but not BMP4/7, and are down-regulated after an early peak in expression. This pattern is exemplified by *tmcc1* in which is expressed non-monotonically in the endoderm trajectory (Fig. 6D). A second pattern described genes that respond to both activin and BMP4/7 with transient non-monotonic expression, such as *nptx2* (Fig. 6E). A third pattern that emerged from this analysis was genes displaying a monotonic increase in expression only to activin, such as *ca14,* suggesting a role in the endoderm lineage specifically (Fig. 6F). Similarly, genes with monotonic and sustained expression in response to BMP4/7, such as *pygm* may be novel mesoderm regulatory factors (Fig. 6G). While these genes are largely unstudied in mesendoderm formation, published transcriptome data sets provide further support for their involvement in mesendoderm formation. For example, *ca14*, *pygm* and *tmcc1* were among genes upregulated in stage 12 animal caps in response to *wnt* and *nodal2* (Ding et al. 2018) and *ca14*, *tmcc1*, and *nptx2* were upregulated in stage 11 embryos in response to somatic cell nuclear transfer from an endoderm cell (Hormanseder et al. 2017). Similarly, p*ygm* has been identified as a target of *myod* (McQueen and Pownall 2017). Thus, using *limma* analysis as an unbiased approach for detecting genes sets that share expression pattern dynamics allows identification of potential new members of developmental GRNs using our data sets.

### Early BMP Signaling Drives Ventral Mesoderm rather than Early Epidermal Divergence

As discussed above, endogenous BMP4/7 signaling within animal pole cells will direct these cells to give rise to epidermis in the absence of BMP inhibitors, which in vivo are secreted by the organizer. BMP signaling is detectable in these cells by stage 10.5, as evidenced by detection of pSmad1/5/8 (Fig. 4A). When explants are treated with exogenous BMP4/7 at stage 9, pSmad1/5/8 is detected at stage 10 at levels comparable to those seen at stage 10.5 in untreated explants (Fig. 7A). This allows a quantitative comparison of the transcriptional responses to the same signal when presented with shifted developmental timing – a tilting of the landscape topology. One predicted outcome of such a heterochronic shift might have been an accelerated transit to the epidermal state rather than formation of ventral mesoderm. To examine how shifting the timing of BMP activity alters the transcriptional response we first compared the transcriptome dynamics. In this context it is interesting to note that premature BMP signaling actually dampened early gene expression changes – there is an ∼32% reduction in the number of genes whose expression significantly changes between stages 9 and 10 (Fig. 7B). This is driven almost entirely by a reduction in the number of genes whose expression decreases during those initial stages (Fig. 7C). Between stages 10 and 11, the transcriptome dynamics are comparable in the two conditions (Fig. 7B,C). However, the number of genes displaying temporal differential expression that are shared by these two trajectories but not by the neural and endodermal state transitions (making them a general response) is remarkably low, ranging from 97 between stages 10 and 10.5, to 55 from stages 10.5 to 11 (Fig. S3 C-F).

**Figure 7.**
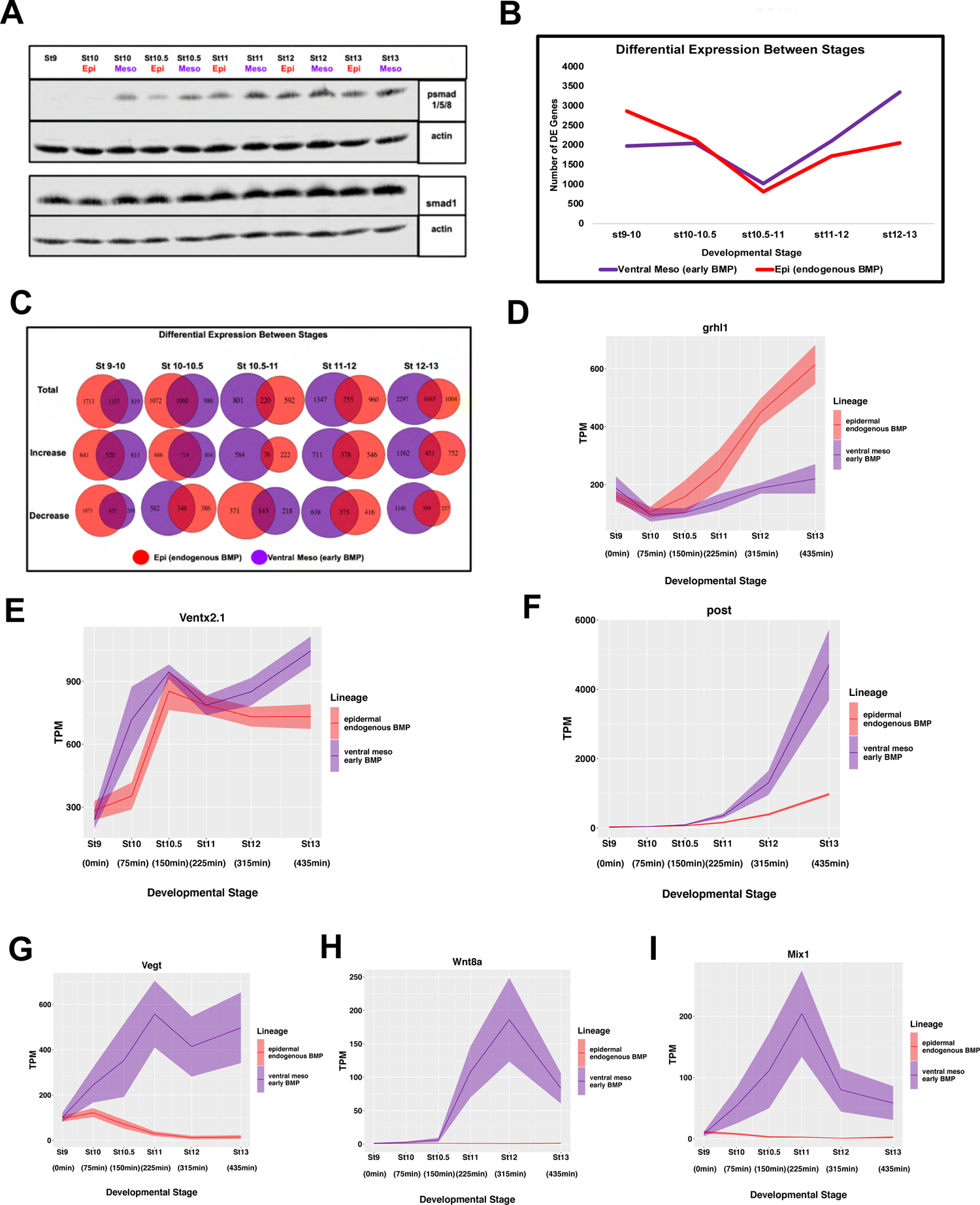
Early BMP signaling drives ventral mesoderm rather than early epidermal divergence. (A) Western Blot Analysis of lysates for developing epidermal (WT) and ventral mesoderm (BMP4/7 20ng/uL) explants for psmad1/5/8 and smad1 with actin loading control. (B) Number of differentially expressed genes between successive developmental stages of the epidermal and ventral mesoderm lineage, *p*_adj_ ≤ 0.05 (C) Venn Diagrams for the total number of DE genes between stages for the epidermal and ventral mesoderm lineages, as well as the number of genes increasing and decreasing over time. (D-I) TPM of genes representative of different expression dynamics in response to early BMP (D) *grhl1* (E) *Ventx2.1* (F) *Post* (G) *Vegt* (H) *Wnt8a* (I) *Mix1*. Graphs are sums of S+L allele. Width of the line represents SEM of three biological replicates.

We next used DESeq2 to compare the genes differentially expressed in response to early BMP signaling. At stage 10 there are only 332 genes differentially expressed between these two conditions. The number of differentially expressed genes remains relatively low until stage 11, when it begins to increase over time (Fig.S6C). However, beginning at stage 10 the genes whose expression is higher in response to early BMP are enriched for mesoderm GO terms (Fig. S6 D). Because expression of *grhl1* is a key early driver of transit to the epidermal state (Tao et al. 2005), we examined its response to early BMP signaling. Strikingly, its expression fails to be robustly activated under this condition and the differential response is seen as early as stage 10.5 (Fig. 7D). Given this surprising finding we examined the expression of *Ventx2.1*, which is a direct target of BMP signaling (Onichtchouk et al. 1996). Here there was a shift in transcription dynamics that reflected the earlier onset of BMP signaling; *Ventx2.1* transcripts reach levels at stage 10 that would not be achieved until state 10.5 in response to endogenous BMP signals (Fig. 7E). Importantly, this demonstrates that BMP signaling is able to immediately elicit changes in gene expression when activated prematurely, and that this drives a heterochronic response in expression of some BMP target genes. Another BMP responsive gene, *Post*, which plays a role in conferring posterior/ventral attributes to both ectoderm and mesoderm (Sato and Sargent 1991) did not show a premature onset of expression, but instead displayed a significant increase in its amplitude of expression (Fig. 7F). Most striking, however, was the activation of genes categorized as activin/nodal target genes including *Vegt*, *Wnt8a*, and *Mix1* (Fig G-I)(Christian et al. 1991, Rosa 1989). Importantly, activation of these genes in response to early BMP is not due to the inappropriate activation of pSmad2/3 (Fig. S5 A). We also confirmed that is the early timing and not the exogenous nature of BMP4/7 exposure that drives ventral mesoderm formation. While treatment of stage 10.5 explants with BMP4/7 leads to increased pSmad1/5/8 (Fig. S6 A), it does not lead to expression of mesodermal genes (Fig. S6 B).

### BMP Signaling is Restrained Until Stage 10.5 by dand5 Activity

The striking finding that shifting the timing of BMP signaling leads to activation of activin/nodal-responsive genes, and the formation of mesoderm instead of epidermis, indicates that it is essential to tightly control when cells receive this signal endogenously. *BMP2,4 and 7* ligands, as well as the receptor s*mad1* are expressed at stage 9 and 10 (Fig. 8A) and cells are clearly competent to respond to early BMP signals as evidenced by the shifted activation of *Ventx2.1* (Fig. 7.E). Despite this, however, pSmad1/5/8 is not robustly detected in control explants until stage 10.5. We therefore investigated how BMP signaling is restrained in animal pole cells until the onset of gastrulation. We asked if there were BMP signaling antagonists expressed in blastula animal pole cells that were downregulated with dynamics consistent with the observed timing of pSmad1/5/8accumulation. We identified two maternally provided BMP antagonists, *dand5* and *gtpbp2* (Bell et al. 2003, Bates et al. 2013, Reich and Weinstein 2019, Kirmizitas et al. 2014), that are expressed at stage 9 but downregulated to significantly lower levels by stage 10.5, when BMP activity is observed (Fig. 8B). *Dand5*, in particular, is robustly expressed at stages 9 and 10 in animal pole cells. To determine if dand5 plays a role in preventing premature BMP signaling we used a translation blocking morpholino to deplete it from early blastulae. We found that dand5-depletion led to premature phosphorylation of smad1/5/8, indicative of early BMP signaling (Fig. 8C), as well as expression of mesodermal marker *xBra (t)* at stage 13 (Fig. 8D). This suggests that a key role for *dand5* at these stages is preventing premature BMP signaling that would generate a mesodermal rather than ectodermal state transition. As *dand5* depletion does not increase pSmad1/5/8 or brachyury levels to the same degree as BMP4/7 treatment, other BMP antagonists including *gtpbp2,* likely cooperate in temporally constraining BMP signaling.

**Figure 8.**
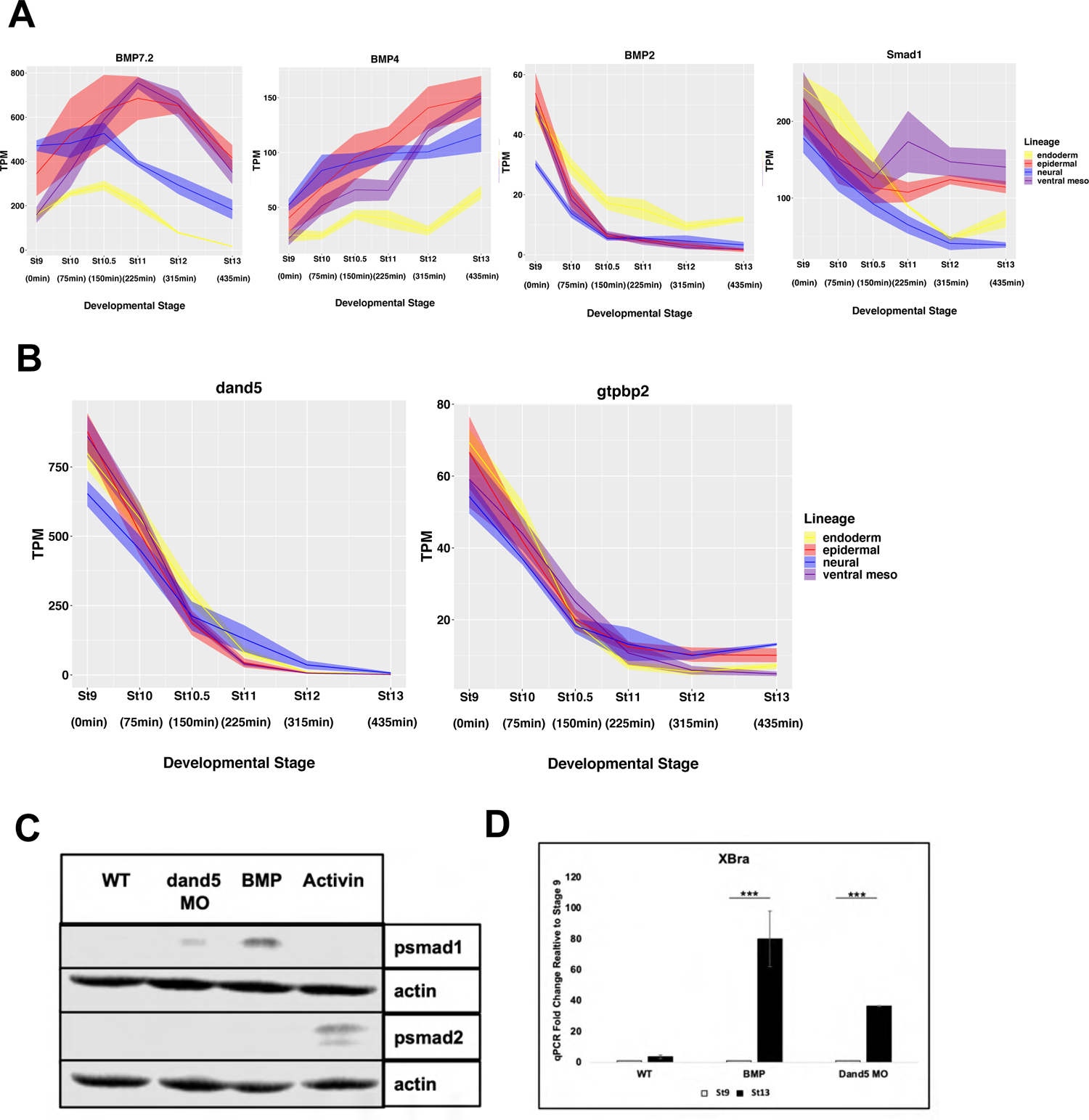
BMP Signaling is Restrained Until stage 10.5 by dand5 Activity. (A-B) TPM of genes involved in BMP signaling (A) BMP heterodimer ligands *BMP7.2, BMP4, BMP2* and primary BMP target smad *smad1* (B) maternally provided BMP antagonists *dand5* and *gtpbp2*. Graphs are sums of S+L allele. Width of the line represents SEM of three biological replicates. (C) Western Blot Analysis of lysates for stage 10 epidermal (WT), *dand5 MO* injected (40pmol/embryo), ventral mesoderm (BMP4/7 20ng/uL), and endoderm (160ng/uL activin) explants for psmad1/5/8 and psmad2 with actin loading control (D) qRT-PCR of animal pole explants examining the fold change from stage 9 to 13 of expression of mesodermal marker *Brachyury(XBra)* for epidermis(WT), ventral mesoderm(BMP4/7 20ng/uL) and *dand5*MO(300pmo/embryo) injected(\*\*\**P*<0.005).

## Discussion

How progenitor cells decide their fate is a fundamental question central to all of developmental biology. While in many cases the inductive cues that drive these decisions have been identified, less is understood about how the timing of these signals is controlled and the dynamics of the transcriptional circuitry they activate. A related and fascinating question is how a large and complex set of transcriptional responses is canalized into a discrete set of lineage trajectories.

Decades of research in *Xenopus* and other systems has shed important light not only on the signaling pathways driving lineage formation in early embryos, but also on many of the key transcriptional targets of these signals. Combined with gain and loss of function studies for individual factors, this work has allowed the construction of putative gene regulatory networks (GRNs) depicting how different lineage states are adopted. A powerful strength of the *Xenopus* system is the ability to easily isolate pluripotent cells from blastula embryos and culture them in simple saline with or without added inductive cues, and that these cells become lineage restricted over a period of approximately seven hours. This allows the transcriptional responses to inductive cues to be quantified with high temporal resolution, enabling the dynamics of individual genes as well as entire lineages to be followed. We used the pipeline we established here to study the transit of initially pluripotent cells to four distinct lineage states using bulk transcriptomics. However, this pipeline can be built upon to layer on additional analyses including ATAC to follow changes in chromatin accessibility and ChIP-seq to follow changes in epigenetic marks dynamically during lineage restriction.

### Neural as the Default State

Our analysis of four different state transitions lends unexpected support for the neural default model, and provides novel insights into why neural is the default state for pluripotent cells. Two types of analysis immediately distinguished the neural lineage from the other three state transitions. First, temporal DESeq reveals that this lineage reaches a steady state in gene expression dynamics by stage 10.5; after this time point the number of genes exhibiting differential expression is quite small (Fig. 2D). This is in marked contrast to the epidermal and mesodermal state transitions which become increasingly dynamic after stage 10.5. Similarly, principal component analysis reveals that the neural state lies closer to the pluripotent state than the other three lineages (Fig. 2B). This is true for both the PC1 and PC2 axes, which together explain 75% of the variation in gene expression. Since developmental times correlate along the PC1 manifold while state identities correlate along the PC2 manifold, this confirms that the time taken to transit from pluripotent to neural state is shorter than for the other lineage trajectories. Measuring the distance between stages 9 and 13 across all 74 principal components also indicates that the neural lineage is most positively correlated with pluripotency (Fig. 2C). Thus, neural progenitor cells occupy a position closest in state space to pluripotent cells relative to the other lineages.

The neural default model has not been without controversy. Studies in avian embryos have challenged the model and suggested that BMP inhibition may not be sufficient for transit to a neural progenitor state (Streit et al., 2000; Linker and Stern, 2004). While work in mESCs and hESCs provided additional support for the neural default model (Smukler et al. 2006, Vallier et al. 2004), these cells do not necessarily recapitulate the in vivo state of inner cell mass cells. Our findings thus provide important validation of this model. Interestingly, when the neural lineage trajectory is compared to that of the epidermal trajectory they are highly correlated through stage 10.5, the onset of gastrulation. This is true with respect to their dynamics, as evidenced by temporal DESeq (Fig 3a), and also supported by the very small number of genes that exhibit differential expression between these states until stage 10.5 (Fig. 3D). The onset of gastrulation can thus be considered a point in time when a group of equipotent cells (neural/epidermal) diverge and either continue changing state to become epidermal or do not continue changing state and become neural.

Significantly, stage 10.5 is when we first detect robust BMP signaling in animal pole cells, as evidenced by pSmad1/5/8 detection (Fig. 4A), and this correlates with the divergence of the epidermal lineage from that of neural (Fig. 3A). It is also the time when we observe a sharp increase in expression of *grhl1*, a key upstream component of the epidermal GRN (Tao et al. 2005), significant down-regulation of the pluripotency factor *foxi2* (Cha et al. 2012) (Fig 3C) and a loss of enrichment for neural GO terms in the epidermal lineage (Fig. S4 A). Neural features are retained and enhanced in the noggin-treated explants, as exemplified by the expression dynamics of transcription factors *Sox3*, *Sox11* and *Zic1* (Fig. 3E-G). By contrast the BMP target genes *Ventx2.1* and *Id3* exhibit expression maxima in the epidermal lineage and minima in the neural lineages around stage 10.5 (Fig. 4 C, D).

Given the distinct non-monotonic dynamics of *Id3* in the epidermal and neural lineages it is tempting to speculate that this inhibitory bHLH factor may be playing a role in suppressing the function of neutralizing factors in the prospective epidermis thus helping to canalize this state transition.

### Surprising overlap in Transcription response to BMP4/7 and Activin

Our data sets allow direct comparison of the transcriptome changes driven by the two different branches of the TGF-beta superfamily, BMP and activin. A striking feature of the activin-driven endoderm trajectory is its distinct dynamics, characterized by a large increase in differentially expressed genes between stage 9 and stage 10.5, the onset of gastrulation (Fig. 5A). This is not seen following treatment with BMP4/7. As expected, the characterized members of the mesendoderm GRN are induced in response to activin and many of these respond rapidly and robustly, but transiently.

Indeed, twenty-one of thirty nine GRN factors examined are induced by stage 10, only 75 minutes after ligand exposure (Fig. 6A,B). pSmad2/3 is also detectable by this time, but is no longer detected by stage 13 indicating that this response too is transient (Fig. S5 A). Among the genes that respond immediately to activin are several members of the Bix/Mix family of transcription factors (Pereira et al. 2012). Somewhat unexpectedly, genes generally associated with ventral fates, including *ventx2.1*, *ventx2.2* and *wnt8,* were also induced by high levels of activin. Induction of these factors occurred at later points in the trajectory, and their expression correlates with the downregulation of Bix/Mix family genes and other transiently responding factors, including *eomes, nodal*, and *vegt.* Going forward it will be of interest to determine if these ventralizing factors play a direct role in downregulating the expression of the endodermal factors that are expressed only transiently. By contrast, *Sox17* responds immediately to activin and its expression increases linearly through stage 13. *Endodermin* also has a linear response to induction, although the increase in its expression does not commence until state 10.5, possibly reflecting a role for Sox17 in its activation. The distinct dynamics of early responding endoderm genes allowed the identification of putative new members of the endoderm GRN using *limma* analysis (Fig. D-G).

Among the most surprising findings emerging from these studies was the activation by BMP4/7 of genes that are generally characterized as activin/nodal targets. Indeed, among the earliest responses to BMP4/7 were Mix/Bix family genes, and similar to their response to activin their induction was transient (Fig. 5C). Other unexpected responding genes included *Sox17*, *eomes* and *gsc*. As Smad2/3 phosphorylation was not observed in response to BMP4/7, this suggests that the BMP R-Smads are capable of activating expression of these activin/nodal targets given a permissive cellular context. In this respect it is worth noting that *Mix1.1* was previously identified in a screen for BMP4 responsive genes (Meade et al. 1996), supporting our current findings. It is intriguing that the genes exhibiting immediate responsiveness to BMP4/7 are dorsal mesendoderm factors, whereas pan and ventral mesoderm genes, including *xbra* (*t*), *wnt8*, *post*, *msx1* and *evx1*, are turned on later in the trajectory. This suggests that the initial response to BMP signaling is “dorsal” as it is for activin, and that more “ventral” attributes are a secondary response.

### The Timing of BMP signaling is Critical for Proper Lineage Segregation

The level of Bmp4/7 signaling utilized in these experiments was selected to match the level of pSmad1/5/8 levels present in untreated explants at stage 10.5 (Fig. 7A). This allows comparison of the response to the same signal and amplitude but with shifted developmental timing. Receiving the same level of BMP signaling at a slightly earlier time point might have been expected to accelerate transit to the epidermal state. Indeed, the shifted onset of *ventx2.1* expression is consistent with an accelerated response (Fig. 7E). However, explants also respond to BMP4/7 exposure at stage 9 by inducing expression of mesendodermal factors, including *mix1*, *vegt* (Fig. 7 G, I), and by suppressing the endogenous BMP-mediated increase in expression of the epidermal regulatory factor *grhl1* (Fig. 7D). Thus, exposure to BMP4/7 at stage 9 elicits a fundamentally different transcriptional response than does exposure at stage 10.5. This was confirmed by treating explants with exogenous BMP4/7 at stage 10.5, which fails to elicit a mesendoderm response (Fig. S6 B).

Together these findings indicate that it is critical to control the timing at which initially pluripotent cells are able to respond to endogenous BMP signaling. While we first detect pSmad1/5/8 at stage 10.5, it is likely that low levels of signaling are initiated by stage 10, as that is when increased expression of *Ventx2.1* and *Id3* is observed (Fig. 4C, D). Thus, cell undergo a fundamental change in competence between stages 9 and 10. Interestingly, expression of BMP inhibitors such as noggin, chordin, follistatin, and cerberus commences in the presumptive organizer region at late blastula stages (Wesseley et al. 2001), indicating that blocking BMP signaling in the marginal zone is also critical at these stages. This raised the question of how BMP signaling is restrained in blastula animal pole cells such that a mesendoderm response is prevented and ectodermal competence is established.

Using our data sets we identified two maternally provided BMP antagonists, dand5 and gtpbp2 that are expressed at stage 9 but are significantly downregulated by stage 10.5 (Fig. 8B). As *dand5* displayed significantly higher levels of expression, we examined the consequences of depleting it from initially pluripotent explants. Morpholino-mediated depletion of dand5 depletion resulted in premature phosphorylation of smad1/5/8 and expression of *xBra (t)* at stage 13 (Fig. 8C, D) (Fig. 8C), indicating that a key role for *dand5* at these stages is preventing premature BMP signaling that would generate a mesodermal rather than ectodermal state transition. Interestingly, a role for dand5 in controlling the spatial response to activin/nodal signaling had previously been suggested, restricting this response to the mesodermal mantle (Bell et al. 2003, Bates et al. 2013, Reich and Weinstein 2019). Our findings suggest a second centrally important role for this TGF-beta antagonist in the temporal control of BMP signaling as animal pole cells exit from pluripotency. Moreover, the data sets described here will facilitate future studies into the temporal control of transcriptional responses to inductive cues across multiple embryonic lineages.

## Materials and Methods

### Embryological Methods

Wild-type *Xenopus laevis* embryos were obtained using standard methods from a daily 2pm fertilization from a single frog and placed into a 14C incubator at 2:45pm until 8:30am. Ectodermal explants were manually dissected at early blastula (stage 8-9) from embryos cultured in 1x Marc’s Modified Ringer’s Solution (MMR) [0.1 M NaCL, 2mM KCl, 1mM MgSO_4_, 2 mM CaCl_2_, 5mM HEPES (pH 7.8), 0.1 mM EDTA] from 8:30am-9:45am and then placed in a 20C incubator until 5pm. Groups of 12-15 explants were collected at 9:45 am, 11am, 12:15pm, 1:30pm, 3pm and 5pm using sister embryos to confirm approximate stages of 9,10, 10.5, 11, 12, and 13 based on Nieuwkoop and Faber (1994) staging. Explants for the neural progenitor lineage were generated using recombinant noggin protein (R&D Systems) at a final concentration of 100ng/mL media supplemented with 0.1% bovine serum albumin (BSA) as a carrier for sequencing experiments, or using 20uM K02288 (Sigma) in 0.1X MMR for Westerns and IF. Endoderm lineage explants were generated using recombinant activin protein (R&D Systems) at a final concentration of 160 ng/mL in 1XMMR supplemented with 0.1% BSA. Mesoderm lineage explants were generated using recombinant BMP4/7 heterodimer protein (R&D Systems) at a final concentration of 20g/mL in 1xMMR supplemented with 0.1% BSA. For morpholino experiments, a previously validated translation-blocking dand5 morpholino (Vonica and Brivanlou, 2007; Gene Tools, sequence: ‘CTGGTGGCCTGGAACAACAGCATGT’) was injected in 4 cells at the eight-cell stage for a total of 40pmol per embryo.

### RNA Isolation, cDNA Synthesis, Sequencing and qRT-PCR

RNA was isolated from blastula explants (12-15 explants) using Trizol (Life Technologies) followed by LiCl precipitation. 1 ug of purified RNA was used as a template for synthesizing cDNA using a High Capacity Reverse Transcription Kit (Life Technologies) Quantitative (q) RT-PCR was performed using SYBR Premix ExTaq 11 (Takara Bio) and detected using the Bio-Rad CFX96 Connect system. Brachyury primers used were Fwd: GAA GCG AAT GTT TCC AGT TC and Rev: ACA TAC TTC CAG CGG TGG TT.

Expression was normalized to ornitihine decarboxylase (ODC) ODC primers used were Fwd: TGA AAA CAT GGG TGC CTA CA and Rev: TGC CAG TGT GGT CTT GAC AT. The fold change was calculated relative to stage nine samples from the same time course experiment. The results show the mean of three independent biological replicates, with error bars depicting the SEM. An unpaired, two-tailed *t*-test was utilized to determine significance. 500 ng of RNA was used for library prep with TruSeq mRNA library prep kit (Illumina) and sequenced using Next Seq 500 Sequencing (epi,neur, endo) or HiSeq4000(meso).

### Western Blot Analysis

Blastula explants (20 explants/sample) were collected for specified stage / lineage and lysed in TNE phospho-lysis buffer [50 mM Tris-HCl (pH 7.4), 150 mM NaCL, 0.5 mM EDTA, and 0.5% Triton X-100, 2mM Sodium Orthovanadate, 20mM Sodium Fluoride, 10mM B-Glycerophosphate, 1 MM Sodium Molybdate dihydrate} supplemented with protease inhibitors [Aprotinin, Leupeptin and phenylmethylsulfonyl fluoride (PMSF)] and a PhosStop phosphatase inhibitor and a complete Mini tablet (Roche). SDS-PAGE and western blot analyses were used to detect proteins and modifications using the following antibodies: anti-phospho SMAD2 (Ser465/467, Sigma, 1:500), Smad2 Polyclonal Antibody (Life Technologies 1:500) Anti-phospho Smad1/Smad5/Smad8 (Ser463/465, Sigma, 1:1000), Smad1 (D59D7, Cell Signal, 1:1000), Actin (A2066, Sigma-Aldrich, 1:5000). For chemiluminescence-based detection, horseradish peroxidase (HRP)-conjugated rabbit secondary antibodies were used (Vector Laboratories, 1:20,000). Results shown are representative of at least three independent experiments.

For the detection and quantification using the Odyssey platform (LI-COR Biosciences), blots were incubated simultaneously with primary antibodies for either psmad1/5/8, smad1 and actin or psmad2, smad2 and actin. Smad phosphorylation was detected using IRDye (LI-COR Biosciences) secondary antibodies and protein amounts were quantified using the Image Studio Lite software (LI-COR Biosciences). Relative smad phosphorylation (psmad1/5/8 or psmad2) was calculated against total smad levels (smad1 or smad2) normalized to actin.

### RNA Seq Processing and Computational Analysis

Read quality was evaluated using FastQC (Andrews 2010). Mapping to *X. laevis* v9.2 genome downloaded from xenbase was performed using RSEM to get TPM values (Li and Dewey 2011). Alignment to X.laevis v9.2 genome was performed using STAR2.6.0 to get raw counts using standard parameters (Dobin et al. 2013). Computational Analysis of RNA Sequencing Data was performed using published R Packages. TPM data is an average of three biological replicates of combined data from S and L alleles for each gene, the width of each line represents SEM and graphs were plotted using ggplot2 (Wickham 2016). Lineage correlation was determined using the ggscatter function with the Pearson correlation method using the ggpubr package (Kassambara 2020). Differential expression analysis was done between successive stages for each lineage and between pairs of lineages at corresponding states using DESeq2 on genes with a minimum raw read count of 15 in any lineage at any stage with significance defined as *p*_adj_ ≤ 0.05 and no fold change cut-off (Love et al, 2014). Overlapping DESeq genes were visualized using VennDiagram and UpsetR (Chen and Boutros 2011, Conway et al 2017). GO and KEGG enrichment was calculated using the GOSeq R package (Young et al. 2012). Top DE genes for heatmaps were determined by ranking based on fold change and only genes with a minimum normalized expression of 10 TPM at the relevant stages were plotted using the pheatmap package (Kolde 2015), expression was depicted using *z*-scores. Principal Component Analysis was done on all genes with a minimum raw read count of 15 in any lineage at any stage with the prcomp function in the default stats package, and visualized using ggfortify (Tang and Horikoshi 2016). The top 2 PCs were determined as the most significant based on the elbow of the PC plot. Distances from stage 9 to 13 were calculated using the dist function. Statistical significance of differences in distance were calculated using the Wilcoxon rank sum, wilcox.test in R. Pattern dynamics were determined for linear, quadratic and cubic patterns with limma analysis using the limma voom R package (Law et al 2014). Potential mesendoderm GRN candidates were selected by examining the 10 genes with most similar differential quadratic and linear dynamics between epidermal and ventral mesoderm. Of these genes, those with minimum expression of 30 TPM, dynamics unique only to the endoderm and/or ventral mesoderm lineages and not already defined as mesendoderm GRN members were identified as possible novel GRN members. Genes graphed were those that could be corroborated with published genomic data sets.

### Vertebrate Animals

All animal procedures were approved by Northwestern University’s Institutional Animal Care and Use Committee, and are in accordance with the National Institutes of Health’s Guide for the care and use of Laboratory Animals.

## Acknowledgements

We thank Joe Nguyen and Rachel Schermerhorm for invaluable technical assistance and the members of the LaBonne laboratory, Madhav Mani, and Eliza Duvall for helpful discussions.

---

**Supplemental Figure 1.**
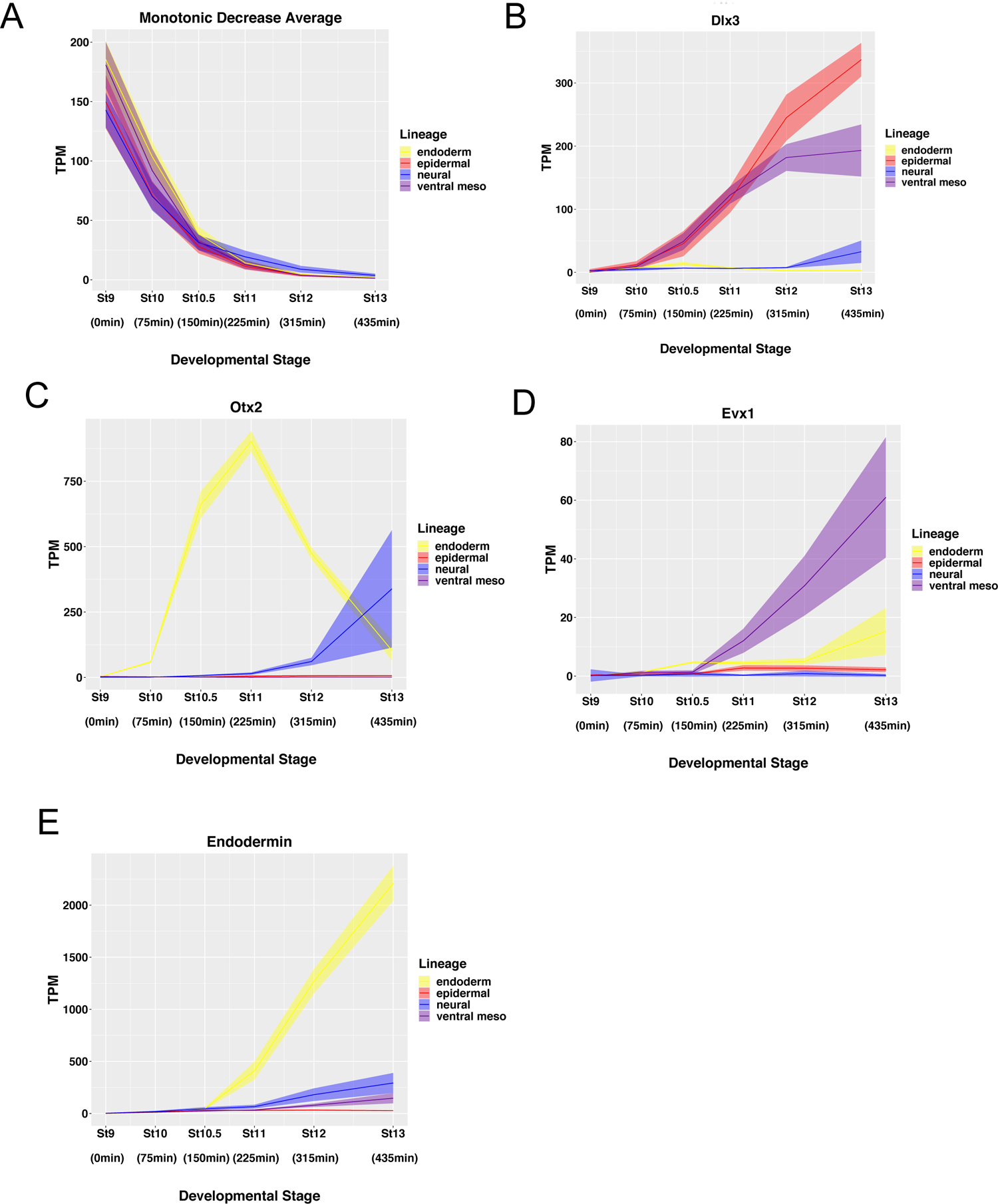
(A-E) RNA Seq TPM expression over time of (A) average of all monotonically decreasing genes with a >25fold decrease between stage 9 and 13 and maximum TPM of 20 at stage 13, (B) epidermal marker *Dlx3*, (C) neural marker *Otx2,* (D) mesoderm marker *Evx1* (E) endoderm marker *Endodermin*. Graphs are sums of S+L allele. Width of lines represents SEM of three biological replicates.

**Supplemental Figure 2.**
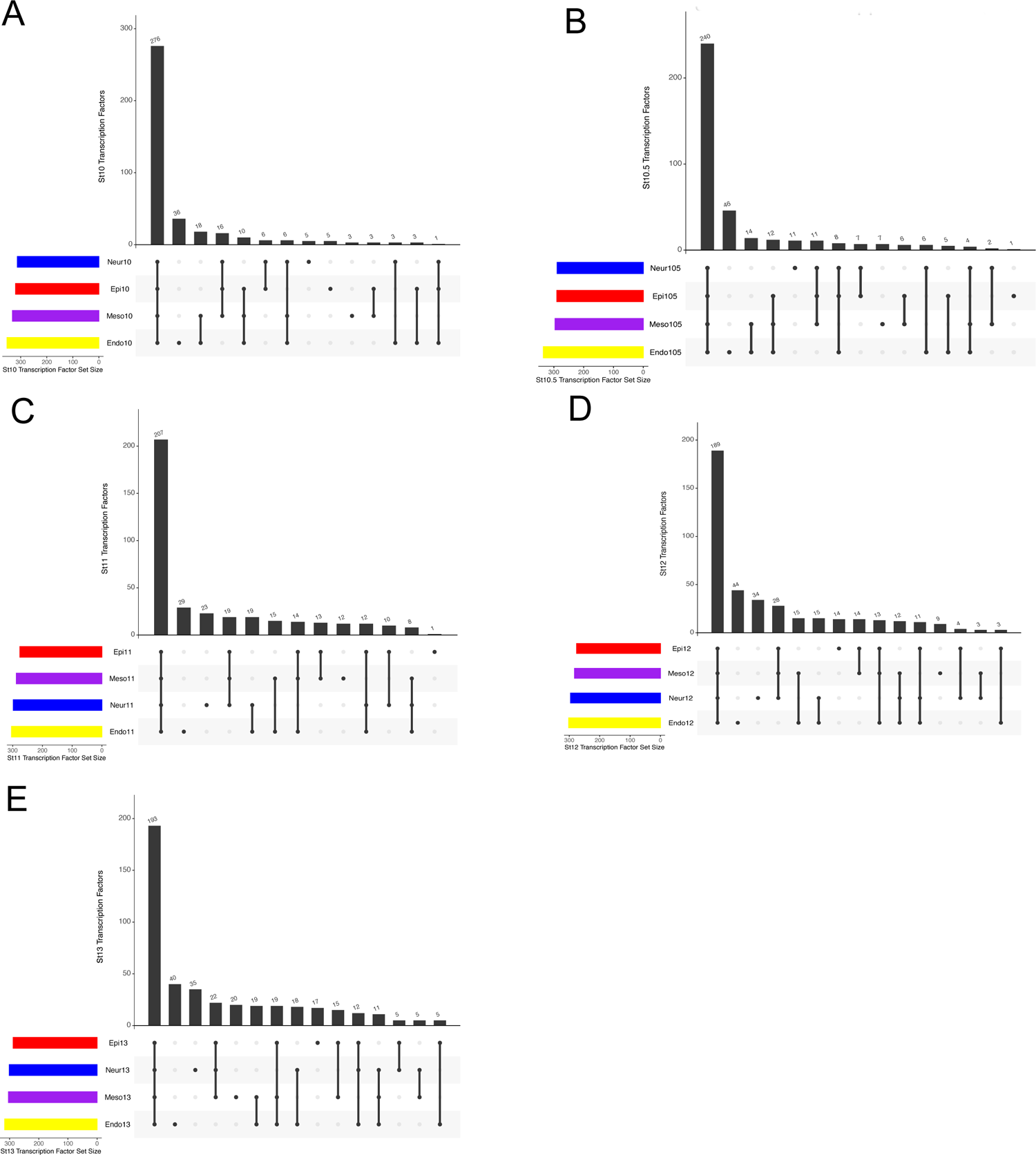
(A-E) UpSet plots of transcription factors expressed at a minimum of 10 TPM at (A) stage 10, (B) stage 10.5, (C) stage 11, (D) stage 12, (E) stage 13. X-axes show genes unique to each lineage and overlapping in all different combinations of lineages ordered from largest number of genes to smallest.

**Supplemental Figure 3.**
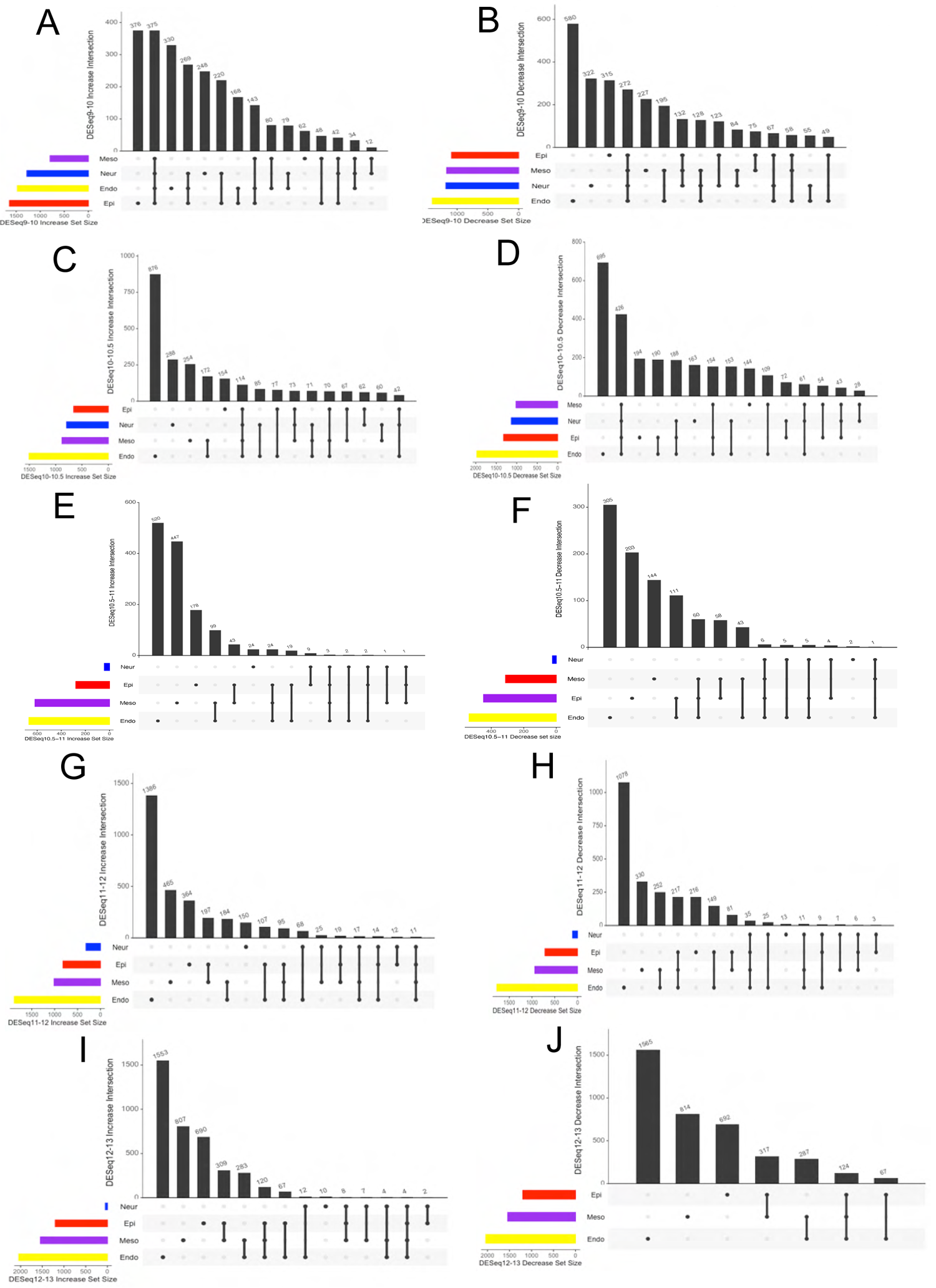
(A-J) UpSet plots of Differentially Expressed Genes (A) increased between stages 9 and 10, (B) decreased between stages 9 and 10, (C) increased between stages 10 and 10.5, (D) decreased between stages 10 and 10.5, (E) increased between stages 10.5 and 11, (F) decreased between stages 10.5 and 11, (G) increased between stages 11 and 12, (H) decreased between stages 11 and 12, (I) increased between stages 12 and 13, (J) decreased between stages 12 and 13.

**Supplemental Figure 4.**
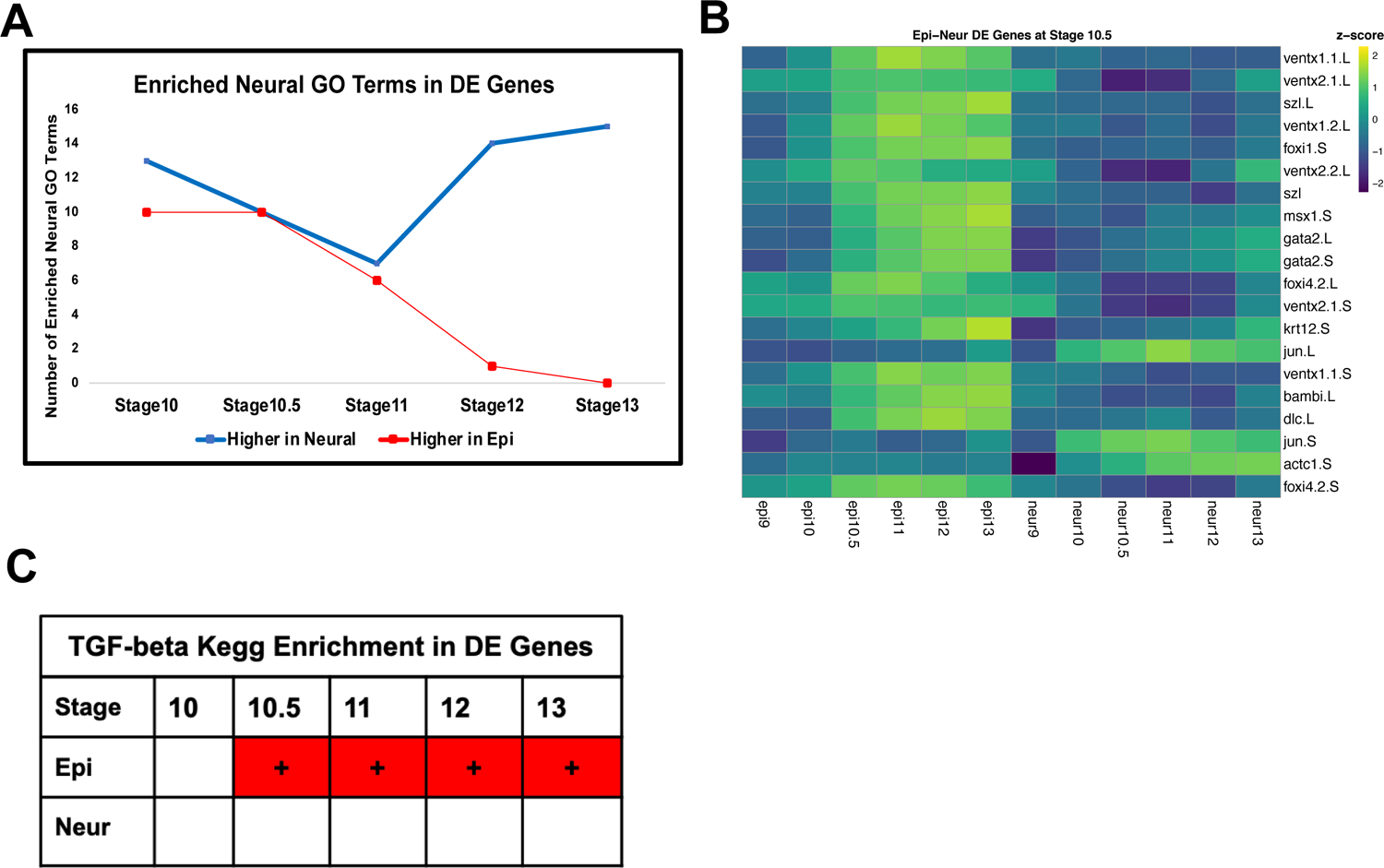
(A) Number of enriched Neural GO Terms in genes significantly higher in the neural lineage (blue) and epidermal lineage (red) at each developmental stage. (B) Heatmap of the top 20 DE genes by Log2FC with a minimum expression of 10TPM between the epidermal and neural lineages at stage 10.5 (C) Kegg enrichment analysis of genes differentially expressed between epidermal and neural lineage at each developmental stage. Genes significantly increased in the epidermal lineage are enriched for TGF-beta genes, as defined by KEGG database from stages 10.5-13.

**Supplemental Figure 5.**
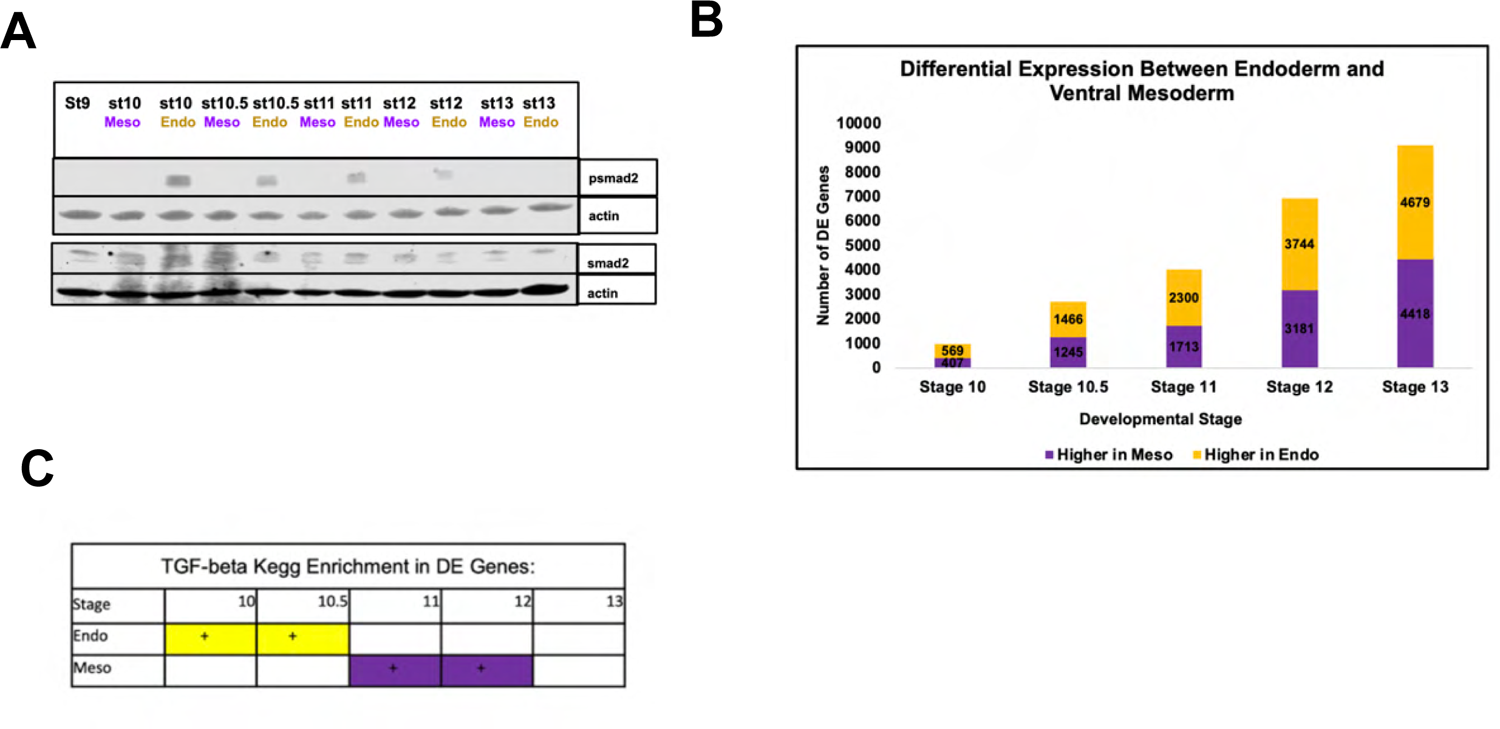
(A)Western blot analysis of lysates of developing mesoderm (20ng/uL BMP4/7) and endoderm (160ng/uL activin) explants for psmad2 and smad2 with actin loading control (B) Number of differentially expressed genes between the endoderm and ventral mesoderm lineages at each developmental stage (*p*_adj_ ≤ 0.05). (C) Kegg enrichment analysis of genes differentially expressed between ventral mesoderm and endoderm lineage at each developmental stage. Genes significantly increased in the endoderm lineage are enriched for TGF-beta genes, as defined by KEGG database from stages 10-10.5 and genes significantly higher in the ventral mesoderm lineage are enriched for TGF-beta genes for stages 11-12.

**Supplemental Figure 6.**
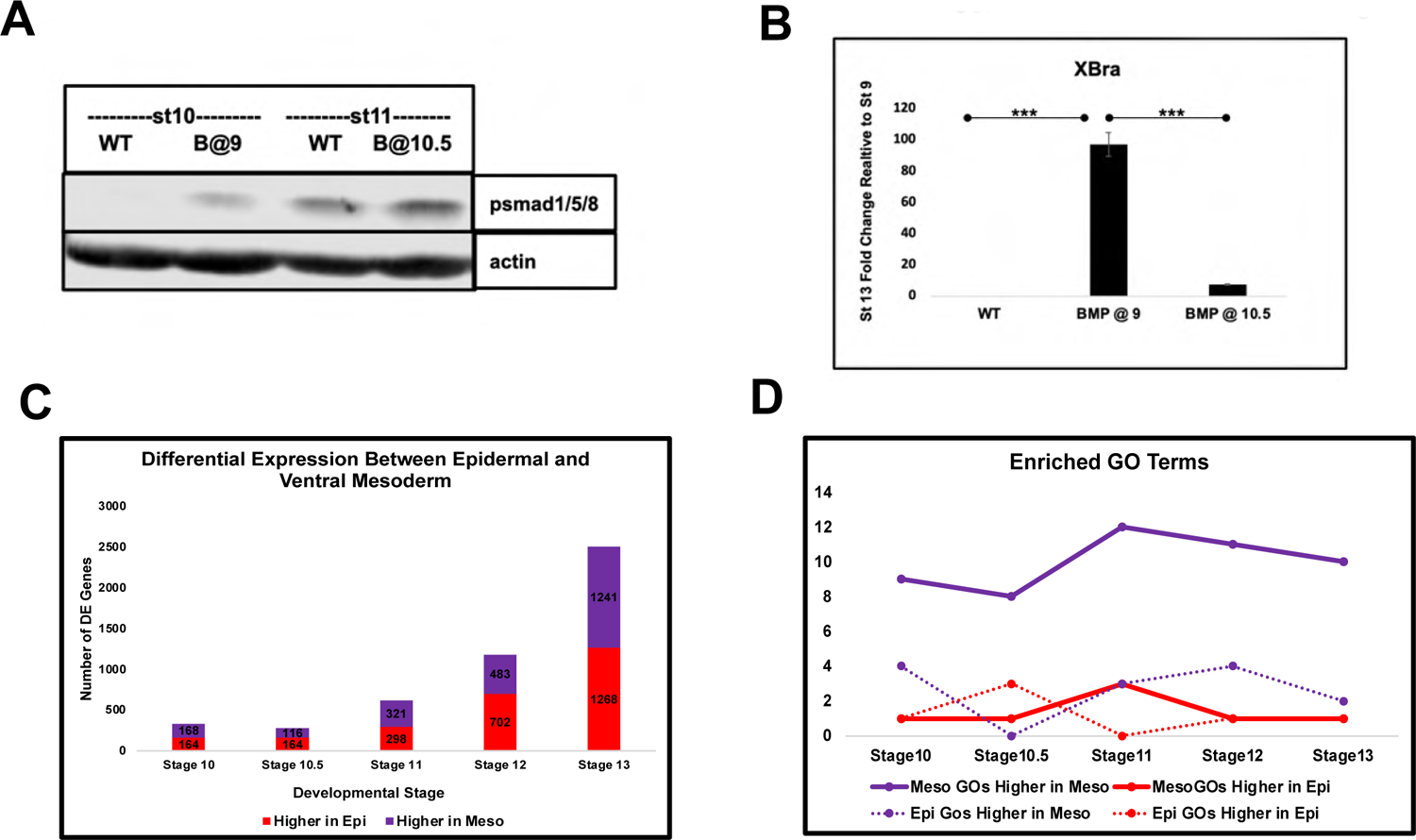
(A) Western Blot Analysis of lysates for epidermal (WT) and BMP4/7 treated at stage 9 (BMP4/7 20ng/uL) explants collected at stage 10 and epidermal (WT) and BMP4/7 treated at stage 10.5 (BMP4/7 20ng/uL) explants collected at stage 11 for psmad1/5/8 with actin loading control (B) qRT-PCR of animal pole explants examining the fold change from stage 9 to 13 of expression of mesodermal marker *Brachyury(XBra)* for epidermis(WT), treated with BMP4/7(20ng/uL) at stage 9 and treated with BMP4/7 (20ng/uL) at stage 10.5 (\*\*\**P*<0.005). (C) Number of differentially expressed genes between the epidermal and ventral mesoderm lineages at each developmental stage (*p*_adj_ ≤ 0.05). (D) Number of enriched Mesoderm GO Terms (solid line) and Epidermis GO Terms (dashed line) in genes significantly higher in the ventral mesoderm lineage (purple) and epidermal lineage (red) at each developmental stage.

